# Prediction of protein-carbohydrate binding sites from protein primary sequence

**DOI:** 10.1101/2024.02.09.579590

**Authors:** Md Muhaiminul Islam Nafi, Quazi Farah Nawar, Tasnim Nishat Islam, M Saifur Rahman

## Abstract

**Background:** A protein is a large complex macromolecule that has a crucial role in performing most of the work in cells and tissues. It is made up of one or more long chains of amino acid residues. Another important biomolecule, after DNA and protein, is carbohydrate. Carbohydrates interact with proteins to run various biological processes. Several biochemical experiments exist to learn the protein-carbohydrate interactions, but they are expensive, time-consuming, and challenging. Therefore, developing computational techniques for effectively predicting protein-carbohydrate binding interactions from protein primary sequence has given rise to a prominent new field of research.

**Result:** In this study, we propose *StackCBEmbed*, an ensemble machine learning model to effectively classify protein-carbohydrate binding interactions at residue level. StackCBEmbed combines traditional sequence-based features along with features derived from a pre-trained transformer-based protein language model. To the best of our knowledge, ours is the first attempt to apply protein language model in predicting protein-carbohydrate binding interactions. StackCBEmbed achieved sensitivity and balanced accuracy scores of 0.730, 0.776 and 0.666, 0.742 in two separate independent test sets. This performance is superior compared to the earlier prediction models benchmarked in the same datasets.

**Conclusion:** We thus hope that StackCBEmbed will discover novel protein-carbohydrate interactions and help advance the related fields of research. StackCBEmbed is freely available as Python scripts at https://github.com/nafiislam/StackCBEmbed.

## 1. Introduction

Nucleic acids, proteins, carbohydrates, and lipids are important biomolecules for any organism. Among these, carbohydrates come right after DNA and proteins in terms of importance [1]. Carbohydrates interact with protein molecules to facilitate various biological processes, such as cellular adhesion, cellular recognition, protein folding, ligand recognition, and so on [2, 3, 4]. Moreover, they provide protection of human cells against pathogens [5] as well as play an important role as biomarkers or drug targets [6, 7, 8, 9]. In order to identify protein-carbohydrate interactions, several experimental techniques have been developed. These include X-ray crystallography, nuclear magnetic resonance (NMR) spectroscopy study, molecular modeling, dual polarization interferometry, and so on. However, weak binding affinity and the synthetic complexity of individual carbohydrates have made these techniques more expensive, time-consuming, and challenging [10]. Therefore, developing a computational technique for effectively predicting protein-carbohydrate binding sites has become an urgent necessity. The computational approaches concentrate on locating the sites of proteins that bind to carbohydrates.

Several computational methods can be found in the literature that leverages structural features to locate protein-carbohydrate binding sites. Taroni et al. [11] first proposed a method that used known protein structure to predict protein-sugar binding sites. They utilized six different attributes of each site, such as solvation potential, residue propensity, hydrophobicity, planarity, protrusion and relative accessible surface area. Through a feature selection process, three of these attributes (residue propensity, protrusion, and relative accessible surface area) were finally selected. These three parameters were then used to calculate the probability of a surface patch being a carbohydrate-binding site. This method provided 65% accuracy for 40 protein-carbohydrate sequences.

A method called COTRAN, proposed by Sujatha and Balaji [12], is able to predict galactose-binding sites from known galactose-binding proteins. Galactose-binding proteins are a special class of proteins, and galactose-binding sites vary across different protein-galactose complexes. No common features are known to be shared by these proteins to specifically bind to their targets. This study assumed that common rules exist for how they recognize galactose. As a result, the study examines 18 protein-galactose 3D structures from 7 different families to identify a unique pattern for galactose binding sites. Each family has similar binding sites, but these sites differ between families.

Another method called InCa-SiteFinder was proposed by Kulharia et al. [13] to predict inositol and carbohydrate binding sites on the protein surface. This method utilizes amino acid propensities and van der Waals interaction energy between protein and a probe. In order to predict glucose binding sites, Nassif et al. [14] utilized several chemical and residue-based features such as charges, hydrophobicity and hydrogen bonding with support vector machine (SVM) as the classifier. Tsai et al. [15] fed probability distributions of interacting atoms in protein surfaces to neural networks and SVMs to predict the binding sites. An energy-based approach was proposed in [16] to identify the protein-carbohydrate binding sites and the residues that have significant roles in binding. This method distinguished 3.3% of residues as binding sites in protein-carbohydrate complexes. In this study, the dominance of Tryptophan (TRP) to communicate with carbohydrates through aromatic-aromatic interactions had also been observed by performing the binding propensity analysis.

Shanmugam et al. proposed a method to identify and study the residues, which are involved in both folding and binding of protein-carbohydrate complexes [17]. They identified the stabilizing residues by means of hydrophobicity, long-range interactions, and conservations. The binding site residues were identified using a distance cutoff of 3.5Å between any heavy atoms in protein and ligand. The residues that were common in stabilizing and binding, are termed key residues. The study found that most of the key residues are present in *β*-strands. All-*α, α* + *β*, and *α/β* structural classes have more key residues than the other protein classes. Nevertheless, the structure-based methods discussed thus far rely on protein structures that are not always available, thereby rendering those ineffective as well as highlighting the importance of sequence-based approaches.

Malik and Ahmed [6] proposed the first sequence-based computational technique in which evolutionary features from the Position Specific Scoring Matrix (PSSM) were used to train a neural network that achieved 87% sensitivity and 23% specificity in a test set of 40 protein-carbohydrate complexes. Pai et al. [18] and Agarwal et al. [19] proposed similar PSSM-based methods to predict mannose-binding sites. Another method, SPRTNT-CBH was proposed in [10] where PSSM features along with other predicted structural features (referred to as sequence-derived structural information) were employed to train an SVM classifier. The classifier achieved 22.3% sensitivity and 98.8% specificity on an independent test set of 50 protein-carbohydrate complexes. Both of the methods mentioned above have issues with imbalanced prediction results, either having high sensitivity and low specificity, or vice versa.

To address these concerns, StackCBPred [20] employed a stacking ensemble classifier to enhance the prediction accuracy in terms of sensitivity (56.5%) and specificity (79.7%). This classifier was trained using a balanced training dataset and incorporated various evolutionary features as well as sequence-based predicted structural features. This model managed to somewhat mitigate the prediction bias observed in previous methods. Canner et al. [21] built two deep learning models, CAPSIF:V and CAPSIF:G, that use structures to predict carbohydrate-protein binding sites. Bibekar et al. introduced PeSTo-Carbs, a geometric transformer model, that predicts carbohydrate-protein binding sites using PDB input. They had proposed two models: PS-G and PS-S. However, there is still potential for further improvement in prediction performance. By incorporating more relevant features and exploring various other machine learning (ML) and deep learning (DL) architectures, the prediction performance could potentially be enhanced further.

In this study, we have introduced a stacking ensemble prediction model named StackCBEmbed. This model seamlessly integrates features directly extracted from protein sequences with attributes derived from a pre-trained transformer-based protein language model. An increasingly successful trend in various protein attribute prediction problems has been the utilization of recently developed protein language models [23, 24]. These pre-trained models enable the generation of informative features for individual proteins, as well as the residues therein, offering enhanced predictive power while being computationally less intensive compared to evolutionary and other features that necessitate time-consuming database searches. To the best of our knowledge, ours is the first work to incorporate protein embedding features from a language model in the context of protein-carbohydrate binding sites prediction. Our method showed a significant improvement in terms of several prediction metrics, outperforming the state-of-the-art carbohydrate binding site prediction methods. The key contributions of our work can be summarized as follows.

- We propose StackCBEmbed, a stacking ensemble prediction model for protein-carbohydrate binding sites. The architecture of the ensemble is determined through an objective approach, searching from a set of base predictors.
- We have investigated features from a transformer-based protein language model (ProtT5-XL-U50) alongside conventional sequence-based and evolutionary features. Our results demonstrate the superiority of the embeddings extracted from the protein language model compared to the other features.
- As the training dataset is heavily imbalanced, we have applied an ensemble random undersampling approach for data balancing.
- In addition to using independent test sets from the literature, we have also prepared an independent test set, named TS46.
- StackCBEmbed outperforms state-of-the-art methods for predicting protein-carbohydrate binding sites in terms of sensitivity and balanced accuracy scores, demonstrating a notable improvement across a number of prediction metrics.

## 2. Materials and Methods

### 2.1. Dataset

Zhao et al. [25] extracted 604 protein-carbohydrate complex structures from PROCARB database [26]. After removing glycoproteins, they selected only those proteins with more than five carbohydrate-binding residues (i.e., residues that have one or more heavy atoms that are within 4.5 Å of any heavy atom of carbohydrates). They further removed the redundant proteins using BLAST-CLUST [27] with a sequence identity cutoff of 30%. They thus obtained 113 protein-carbohydrate complexes. Taherzadeh et al. [10] used the Glyco3d database (http://glyco3d.cermav.cnrs.fr/home.php) to obtain more than 1000 three-dimensional structures of protein-carbohydrate complexes. After combining these two sets, they removed low-resolution structures (*>*3 Å) and homologous proteins (30% sequence identity cut-off). They thus obtained 152 protein-carbohydrate complexes. Out of these, 50 proteins were randomly chosen and separated for the independent test set (TS50), and the remaining set (102 proteins), labeled as TR102, was used for model training in their study. Gattani et al. [20] discarded 2 sequences from TR102, and 1 sequence from TS50, as those contained non-standard amino acids. They referred to the updated sets as the benchmark dataset and TS49, respectively. In our study we have used the benchmark dataset for training our prediction model and TS49 for independent testing. However, we had to leave out two more sequences (IDs: 2c3hB and 2vngB) from the benchmark dataset due to a discrepancy in the number of residues in the FASTA files vs. the PSSM files that we generated thereof.

Gattani et al. created a second independent test set from PROCARB as follows. From the PROCARB604 dataset, the proteins whose IDs matched the protein IDs that were present in the benchmark or TS49 dataset were removed. Redundant proteins with a sequence identity cutoff of 30% were then removed. They thus obtained a set of 88 sequences, which was named TS88. Like them, we too have used this as a second independent test set in our study.

The benchmark dataset contains 98 sequences with 1014 binding residues and 25705 non-binding residues. We randomly selected 30% of these residues to create a validation set, consisting of 275 binding residues and 7741 non-binding residues. The remaining portion was designated as the training set.

Additionally, in this work, we developed a third independent dataset from UniProt (https://www.uniprot.org/ to further demonstrate the efficacy of our proposed method. To create this dataset, we used the advanced search option of UniProt with the following query:

`(ft_binding_exp:”CHEBI:16646”) AND (reviewed:true) NOT (ft_carbohyd:n-linked) NOT (ft_carbohyd:o-linked) NOT (ft_carbohyd:c-linked)`

Our search resulted in 122 proteins. After removing the proteins with no positive sites, 115 proteins remained. We then used BLAST-CLUST [27] with 30% sequence identity, which reduced the dataset to 80 proteins. We further filtered the dataset to include only the proteins that possessed more than five carbohydrate-binding residues and enforced 30% sequence identity cutoff with the benchmark dataset using BLAST-CLUST. The final count of proteins was 46, with 472 positive sites and 18820 negative sites. We named this dataset TS46.

The binding and non-binding residue distributions of the datasets used in our study are listed in Table 1, from which it is evident that all the datasets are highly imbalanced.

**Table 1:** Number of binding and non-binding sites in the datasets.

| Dataset Type | Count Of |  |  |
| --- | --- | --- | --- |
|  | Proteins | Binding sites | Non-binding sites |
| Training set | 98 | 739 | 17964 |
| Validation set |  | 275 | 7741 |
| TS49 | 49 | 508 | 13230 |
| TS88 | 88 | 688 | 26453 |
| TS46 | 46 | 472 | 18820 |

### 2.2. Feature Extraction

Feature extraction is a very important step in the prediction pipeline. In this study, we utilized two types of features: the traditional sequence-based features and the embeddings derived from pre-trained transformer-based protein language models. These features are described below. Notably, after featurization, Yeo–Johnson transformation [28] was applied, followed by standardization, using *PowerTransformer* from *scikit-learn* python package.

#### i. Position specific scoring matrix (PSSM)

Position specific scoring matrix (PSSM) expresses the evolution-derived information in protein sequences. Evolutionarily conserved residues have occupied a prominent place in several biological analyses [29, 30, 31, 32, 33, 34] and they have considerably important functional roles in binding. We generated PSSM using PSI-BLAST [35] with E-value = 0.001, with 5 iterations. The PSSM is a *L ×* 20 dimensional matrix (*L* is the length of the protein sequence) which can be represented by :

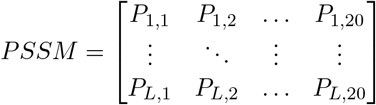

Here, *P*_*i,j*_ indicates the score of the amino acid at position *i* being changed to amino acid *j* during the evolutionary process. For each residue, we thus get a 20-dimensional feature vector.

#### ii. Monogram (MG)

Monogram (MG) [36, 37] feature for a residue is extracted from PSSM by summing up the scores for that residue along the protein sequence’s length. Thus, it results in a feature vector of length 1 for each residue and is location invariant.

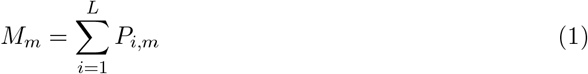

where, 1 *<*= *m <*= 20 and L is the length of the sequence.

#### iii Di-peptide composition (DPC)

Di-peptide composition (DPC) [38] can also be computed from PSSM to capture the partial sequence-order information. As there are 20 *×* 20 combinations of the dipeptides, this feature can be represented as a 20 *×* 20 matrix *DPC*[*i, j*], 1 ≤ *i, j* ≤ 20, which can later be flattened into a 400-dimensional vector. For the residue at position *k*, 1 ≤ *k* ≤ *L, DPC*[*i, j*] is calculated as follows.

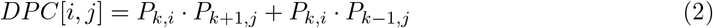

Note that for the first and last residues, the indices *k*− 1 and *k* + 1 respectively are out of range, therefore the corresponding *P* values are assumed to be 0. In other words, for the terminal residues, only one of the two terms in the sum has a contribution.

#### iv Accessible Surface Area (ASA)

In order to predict the structural and functional features of proteins, as well as the sequence-structure-function relationship, the knowledge of solvent exposure is important. Accessible Surface Area (ASA) [39] measures the solvent accessibility of a biomolecule and can play a crucial role in predicting binding sites. Here SPIDER3 program [40, 41, 42] was used to predict ASA probabilities at the residue level directly from each of the protein sequences.

#### v Half Sphere Exposure (HSE)

ASA has several drawbacks, however. For example, it is quite difficult to determine to what extent a residue is buried, and also, it is impossible to differentiate between the partially buried residue and the deeply buried residue through ASA. In order to solve these issues, HSE had been introduced [43]. HSE separates a residue’s sphere into two half-spheres: HSE-up and HSE-down. This measure has several interesting properties in the characterization of the residue’s spatial neighborhood, and it does not rely on a full-atom model. Thus, it is very convenient to apply this measure in protein structure modeling and prediction analysis. HSE-up and HSE-down were also extracted using the SPIDER3 program.

#### vi Torsion angles (TA)

The torsion angles Phi (*ϕ*) and Psi (*ψ*) determine the shape of the protein backbone structure. The predicted values of these two angles have been used as features, as they have crucial roles in analyzing protein-carbohydrate interactions. Here, too, we have used SPIDER3 for the predicted torsion angles.

#### vii Physicochemical properties (PHYSICO)

For each of the 20 standard amino acids, we utilized seven different physicochemical properties, as utilized in [44]: steric parameter, polarizability, normalized Van der Waals volume, hydrophobicity, isoelectric point, helix probability, and sheet probability.

#### viii Embedding from pre-trained protein language model (EPLM)

From the latest Natural Language Processing (NLP) research [24], transformer models were trained on large protein corpora, and these are known as protein language models. The output from the last layer of these models can be used as features in various prediction problems. In our study, we have used ESM2 [45] and six of the models from [24]: ProtBert, ProtBert-BFD, ProtT5-XL-BFD, ProtT5-XL-U50, ProtAlbert, and ProtXLNet. ESM2 is a transformer-based protein language model that is trained over millions of diverse natural proteins across evolution using masked language modeling. Here we have used the esm2 t33 650M UR50D model. ProtBert and ProtBert-BFD are BERT-based models [46] trained on UniRef100 and BFD-100 corpora, respectively. The ProtAlbert model was acquired from the Albert model [47] while ProtXLNet was acquired from the XLNet model [48]. Both were trained on the UniRef100 corpora. On the other hand, ProtT5-XL-BFD was obtained from the T5 model [49] trained on BFD corpora. It was then fine-tuned on the UniRef50 dataset to create the ProtT5-XL-U50 model. Upon comparative analysis of the embeddings for our problem (see Results section), we decided to go with the ProtT5-XL-U50 embedding, which is a 1024-dimensional vector.

### 2.3. Performance Evaluation Criteria

We have used several well-established metrics [50, 51] to objectively measure the performance of our prediction model. These are sensitivity (SN), specificity (SP), accuracy (Acc), balanced accuracy (BACC), Precision (Prec), F1-score (F1), and Matthew’s Correlation Coefficient (MCC). These can be mathematically defined as follows:

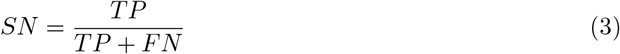

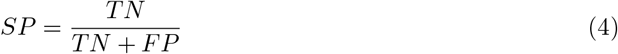

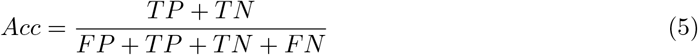

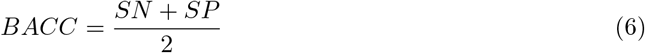

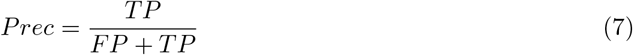

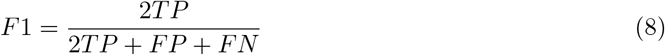

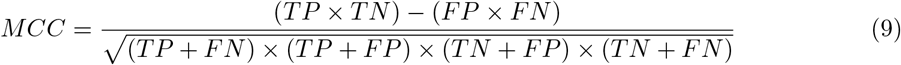

Here, TP, FP, TN, and FN stand for True Positive, False Positive, True Negative, and False Negative, respectively. Additionally, we also assessed the area under receiver operating characteristic curve (AUROC or AUC in short) [52] and area under precision-recall curve (auPR in short) [53].

### 2.4. Feature selection

Using the aforementioned feature descriptors, we used incremental feature selection (IFS) to obtain the optimal combination of features. We took a phased approach to IFS (Algorithm 1), where at each phase, we try to augment the current set of features with one additional feature group. Among all the groups, the augmentation that yields the best performance is chosen, and the feature set is updated. The updated feature set is used in the next phas,e and the process repeats. Thus, while the search space of feature group combinations is exponential, only a limited set of combinations is explored in a greedy manner. Finally, of all the combinations explored, the one with the best performance is chosen. To measure the performance of each combination, we trained a classification model with those features using Extreme Gradient Boosting (XGB) from the scikit-learn package in Python [54], with default parameters. The benchmark dataset, balanced using random undersampling, was used. We used XGB since it has been very successful in numerous ML challenges in Kaggle (https://www.kaggle.com/).

### 2.5. Framework of proposed method

In order to improve the prediction performance, we implemented a stacking ensemble model [55] using two stages of learners, known as *base learners* and *meta learners* respectively. In the first step, multiple base learners were trained using the balanced training dataset. Each trained model was used to make predictions on the validation set. In the next step, the validation set was used to train the meta learner. In this case, however, the feature vector was augmented by concatenating the predictions earlier made by the base learners. These predictions thus form yet another feature descriptor for the meta learner, which we refer to as *base learner predictions (BLP)*.

We experimented with ten machine learning algorithms: Naive Bayes (NB), *K*-nearest Neighbor (KNN), Logistic Regression (LR), Partial Least Square Regression (PLS), Support Vector Machine (SVM), Multi-layer Perceptron (MLP), Decision Tree (DT), Random Forest (RF), Extra Tree (ET) and Extreme Gradient Boosting (XGB). The *xgboost* Python library was used for XGB, while the *scikit-learn* library was used for the remaining learners. For all the learners, the class discriminating threshold was set at the default value, which is 0.5.

We analyzed the mutual information (MI) [56] between the residue-wise prediction probabilities and the target variable for each of these learners. As SVM had the highest MI, it was added in

#### Algorithm 1: Incremental Feature Selection

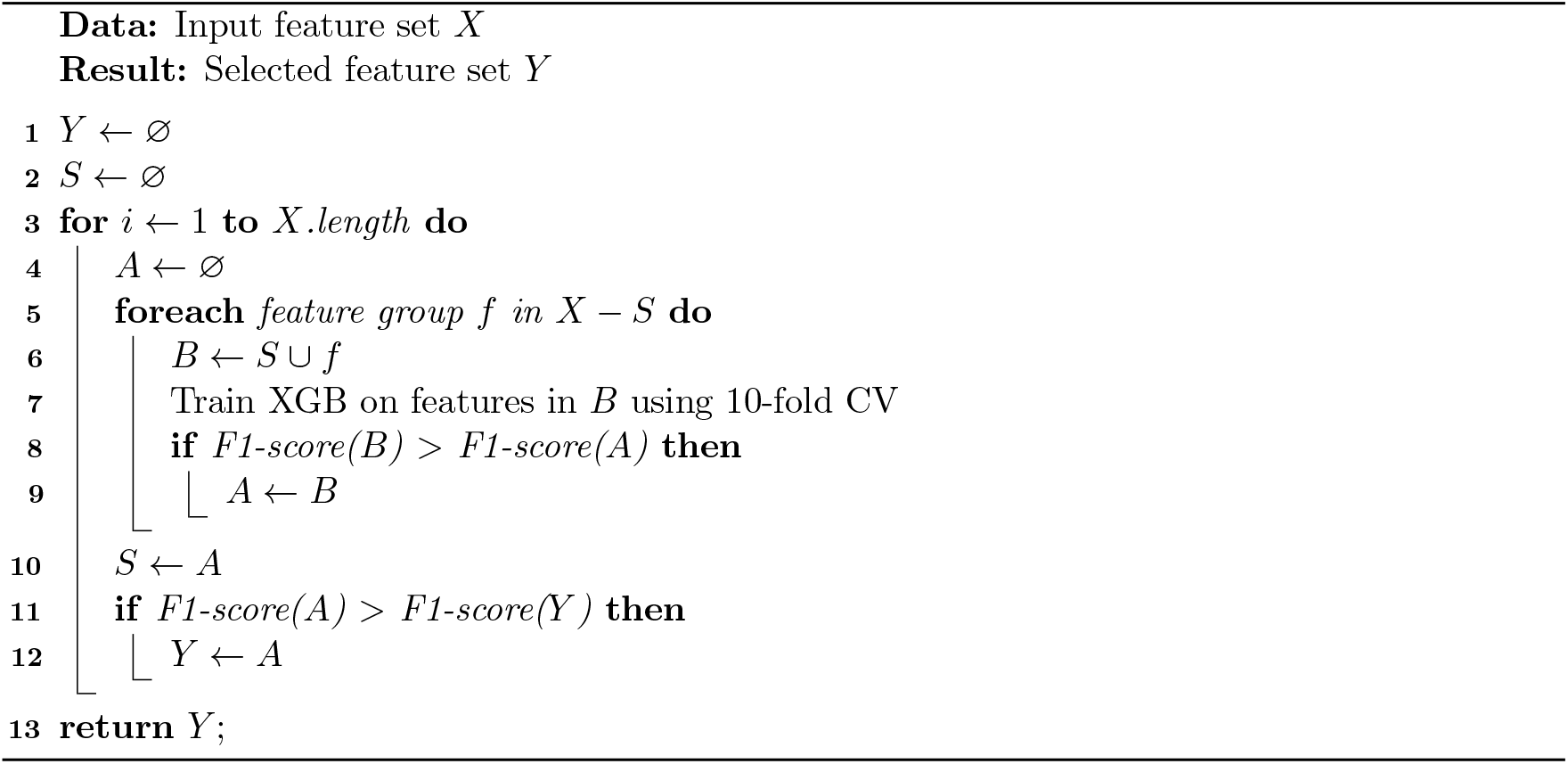

both the base and meta layers. Subsequently, more predictors were added to the base layer in a phased approach as follows. At each phase, we tried to augment the current set of predictors with another predictor and calculated the MI between average prediction probabilities and labels. The predictor for which the average MI became the highest in the phase was added. It is possible that the MI may decrease in a phase from an earlier phase. Nevertheless, the updated set of predictors is used in the next phase, and the process repeats. Finally, the set of predictors that resulted in the highest MI across all the phases is chosen. Accordingly, four classifiers, SVM, MLP, ET, and PLS were selected as the base learners (see Results Section).

### 2.6. Imbalanced data handling

The datasets used in this study were significantly imbalanced. Therefore, it was necessary to balance the training dataset. We applied an ensemble random undersampling approach, inspired by [57], as follows. We prepared 10 subsampled datasets, each of which contained all positive samples. As to the negative samples, four of these datasets contained 0.2% of the negative samples, randomly selected, three datasets contained 2%, and the remaining three datasets contained randomly selected 20% of the negative samples. Using each of these 10 subsampled datasets, we trained separate SVM, MLP, ET, and PLS models. The validation set was featurized using selected feature descriptors, then augmented with the output of the base layer models, which we have referred to as the BLP descriptor. It was then used to train the SVM model in the meta layer. Data balancing in the meta layer was done using random undersampling.

### 2.7. Hyperparameter tuning

We tuned the hyperparameters of the base and meta layer classifiers to optimize prediction performance. We only tuned the crucial hyperparameters. The rest were set at their default values. We used *GridSearchCV* from the *scikit-learn* package in Python to grid search the hyperparameter space. We selected the values of each selected parameter of each classifier that had balanced performance across all performance metrics. After tuning the base classifiers, we applied the same process to the meta layer classifier on the validation dataset.

## 3. Results

### 3.1. Embedding from T5 model outperforms other embeddings

The 10-fold CV results averaged across all 10 predictors trained on protein embeddings from the seven different pre-trained language models are shown in Table 2. ProtT5-XL-U50 embedding wins with respect to all metrics but sensitivity. However, the difference between it and ESM2 in the sensitivity metric is negligible. The ESM2 came out second best in terms of the other metrics. Therefore we selected the ProtT5-XL-U50 embedding for our model.

**Table 2:** 10-fold CV performance of embeddings from various pre-trained protein language models. Results are averaged across models trained using various learning algorithms on the benchmark dataset balanced with random undersampling.

| Language Model | SN | SP | BACC | Acc | Prec | F1 | MCC | AUC | auPR |
| --- | --- | --- | --- | --- | --- | --- | --- | --- | --- |
| ProtBert | 0.697 | 0.719 | 0.708 | 0.708 | 0.715 | 0.704 | 0.418 | 0.775 | 0.766 |
| ProtBert-BFD | 0.712 | 0.718 | 0.715 | 0.715 | 0.717 | 0.714 | 0.431 | 0.780 | 0.773 |
| ProtAlbert | 0.726 | 0.726 | 0.726 | 0.726 | 0.728 | 0.726 | 0.454 | 0.792 | 0.784 |
| ProtXLNet | 0.701 | 0.579 | 0.640 | 0.640 | 0.630 | 0.661 | 0.284 | 0.682 | 0.650 |
| ProtT5-XL-BFD | 0.730 | 0.772 | 0.751 | 0.751 | 0.765 | 0.745 | 0.505 | 0.819 | 0.811 |
| ProtT5-XL-U50 | 0.738 | **0.787** | **0.762** | **0.762** | **0.779** | **0.756** | **0.528** | **0.830** | **0.825** |
| ESM2 | **0.739** | 0.779 | 0.759 | 0.759 | 0.772 | 0.754 | 0.521 | 0.824 | 0.82 |

### 3.2. Embedding from language model wins over traditional feature descriptors

We wanted to examine how each feature descriptor influences the prediction of protein-carbohydrate binding sites. For each feature descriptor, we trained ten ML algorithms as stated before - NB, KNN, LR, PLS, SVM, MLP, DT, RF, ET, and XGB. The averages across these classifiers were used to compare the predictive ability of the descriptors. As observed from Table 3, the embedding from ProtT5-XL-U50 exhibited the highest average scores across all performance metrics, indicating its effectiveness in predicting protein-carbohydrate binding sites. DPC is the best descriptor among the traditional descriptors, followed by PSSM.

**Table 3:**
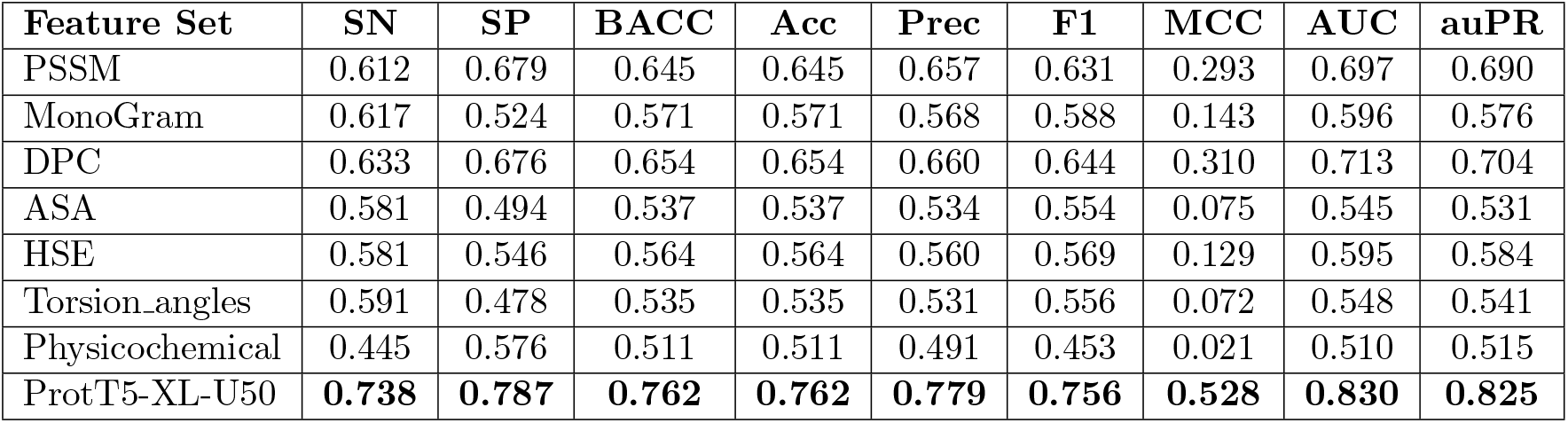
10-fold CV performance of various feature descriptors. Results are averaged across models trained using various learning algorithms on the benchmark dataset balanced with random undersampling.

### 3.3. Protein embedding along with PSSM constitutes the optimal feature descriptor

We had seven traditional feature descriptors and one embedding feature descriptor (ProtT5-XL-U50) at our disposal. As described earlier, to identify the optimal subset of features from this set, we employed incremental feature selection (Algorithm 1). Supplementary Table S1 shows the results of adding different feature groups. Based on the F1-score, we find the combination of ProtT5-XL-U50 embedding and PSSM to be the optimal feature descriptor.

### 3.4. SVM, MLP, ET, and PLS constitute the base layer of StackCBEmbed

Table 4 reports the highest MI obtained in each phase and the corresponding learner combination. When multiple combinations in a phase resulted in very close MI values, all combinations were considered for exploration in the next phase. For example, in phase 3, {SVM, MLP, RF} and {SVM, MLT, ET} produced very close MI values. So we expanded the search in phase 4 based on both these combinations. Based on the overall results, SVM, MLP, ET, and PLS were selected as the base learners. The results from the full search for base layer learners is available in [Supplementary file](https://docs.google.com/spreadsheets/d/1slN-Yyfr4Em0h-aN-aXX92JWoLg_G_yETnmF-1A5L_k/edit#gid=2018168915).

**Table 4:** Mutual information (MI) between the predictions and labels for the different subsets of predictors. For each subset, the probability of binding sites is averaged across the predictors. All the predictors were trained on the training dataset balanced with random undersampling.

| Phase | Best Learner Combination | MI |
| --- | --- | --- |
| 1 | SVM | 0.0430 |
| 2 | SVM, MLP | 0.0495 |
| 3a | SVM, MLP, RF | 0.0492 |
| 4a | SVM, MLP, RF, KNN | 0.0497 |
| 5a | SVM, MLP, RF, KNN, XGB | 0.0511 |
| 6a | SVM, MLP, RF, KNN, XGB, ET | 0.0516 |
| 7a | SVM, MLP, RF, KNN, XGB, ET, NB | 0.0487 |
| 8ab | SVM, MLP, RF, KNN, XGB, ET, NB, LR | 0.0445 |
| 8ac | SVM, MLP, RF, KNN, XGB, ET, NB, PLS | 0.0445 |
| 9a | SVM, MLP, RF, KNN, XGB, ET, NB, LR, PLS | 0.0441 |
| 10a | SVM, MLP, RF, KNN, XGB, ET, NB, PLS, LR, DT | 0.0434 |
| 3b | SVM, MLP, ET | 0.0491 |
| **4b** | **SVM, MLP, ET, PLS** | **0.0521** |
| 5b | SVM, MLP, ET, PLS, RF | 0.0513 |
| 6b | SVM, MLP, ET, PLS, RF, KNN | 0.0488 |
| 7b | SVM, MLP, ET, PLS, RF, KNN, LR | 0.0495 |
| 8b | SVM, MLP, ET, PLS, RF, KNN, LR, NB | 0.0498 |
| 9b | SVM, MLP, ET, PLS, RF, KNN, LR, NB, DT | 0.0452 |

### 3.5. SVM is chosen as the Meta layer classifier

As SVM had the highest MI score (Table 4), it was also chosen as the meta layer classifier. Since the meta layer will be trained on the validation set, we further evaluated all the classifiers using 10-fold CV, and the results are shown in Table 5. SVM has the most balanced performance. Therefore, our choice of SVM for the meta layer has been well-justified.

**Table 5:** 10-fold CV performance of models trained on the validation set using embedding and PSSM features. The validation set was balanced using random undersampling.

| Classifier | SN | SP | BACC | Acc | Prec | F1 | MCC | AUC | auPR |
| --- | --- | --- | --- | --- | --- | --- | --- | --- | --- |
| RF | 0.775 | **0.829** | 0.802 | 0.802 | 0.822 | 0.795 | 0.609 | 0.872 | 0.883 |
| ET | 0.786 | 0.822 | 0.804 | 0.804 | 0.819 | 0.799 | 0.611 | **0.881** | **0.889** |
| DT | 0.691 | 0.695 | 0.693 | 0.693 | 0.695 | 0.692 | 0.387 | 0.693 | 0.636 |
| MLP | **0.840** | 0.692 | 0.766 | 0.765 | 0.734 | 0.782 | 0.540 | 0.837 | 0.845 |
| LR | 0.742 | 0.732 | 0.737 | 0.736 | 0.738 | 0.737 | 0.476 | 0.817 | 0.824 |
| SVM | 0.815 | 0.801 | **0.808** | **0.807** | 0.807 | **0.809** | 0.618 | 0.877 | 0.884 |
| NB | 0.767 | 0.811 | 0.789 | 0.789 | 0.803 | 0.782 | 0.582 | 0.836 | 0.796 |
| KNN | 0.720 | 0.790 | 0.755 | 0.755 | 0.776 | 0.744 | 0.514 | 0.807 | 0.790 |
| XGB | 0.782 | **0.829** | 0.806 | 0.805 | **0.826** | 0.797 | **0.619** | 0.873 | 0.875 |
| PLS | 0.793 | 0.794 | 0.793 | 0.793 | 0.800 | 0.793 | 0.590 | 0.867 | 0.876 |

Figure 1 captures the overall training process of StackCBEmbed. The workflow for predicting the binding sites in a query protein is demonstrated in Figure 2.

**Figure 1.**
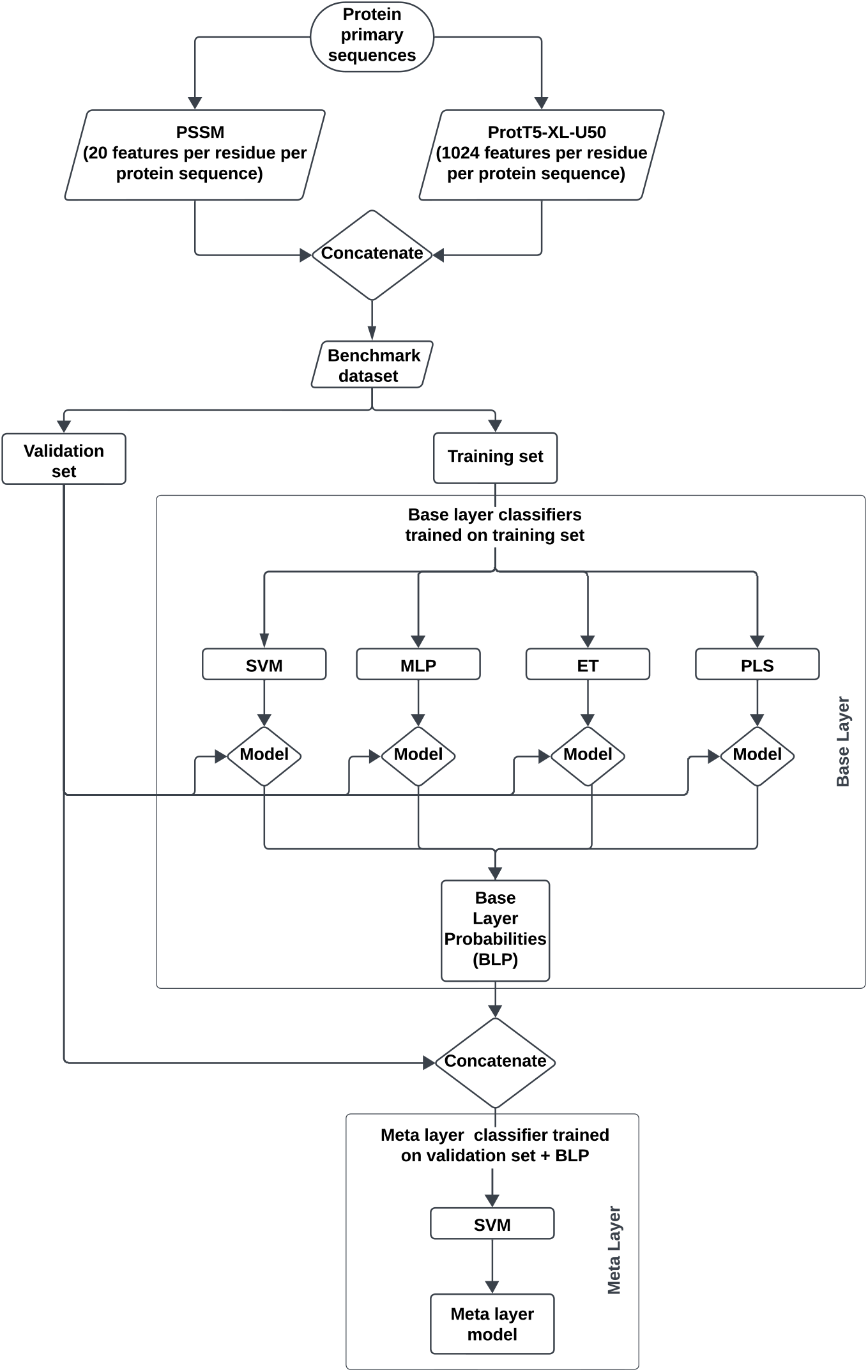
The framework and training process of StackCBEmbed.

**Figure 2.**
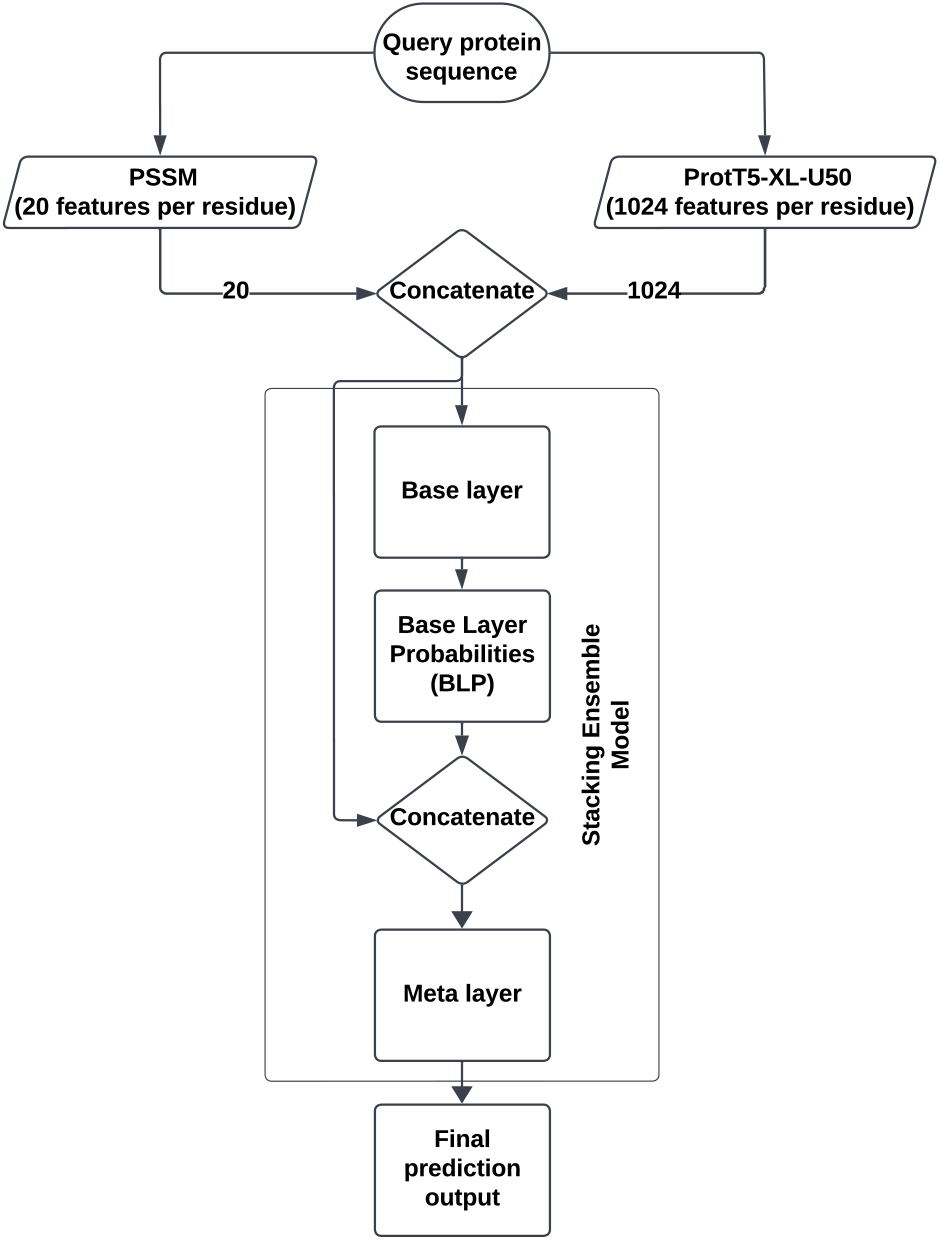
StackCBEmbed workflow for protein-carbohydrate binding-site prediction for a novel/query protein.

### 3.6. Hyperparameter tuning

The hyperparameters for the base layer classifiers (SVM, MLP, ET, PLS) and the meta layer classifier (SVM) were tuned using *GridSearchCV*. The parameters we tuned, alongside their final values, are given in Table 6. (See [Supplementary file](https://docs.google.com/spreadsheets/d/1slN-Yyfr4Em0h-aN-aXX92JWoLg_G_yETnmF-1A5L_k/edit?gid=1118520371#gid=1118520371) for the full grid search results).

**Table 6:** List of hyperparameters with their tuned values.

| Layer | Classifier | Parameters | Values |
| --- | --- | --- | --- |
| Base layer | SVM | C | 10 |
|  |  | degree | 3 |
|  |  | gamma | auto |
|  |  | kernel | rbf |
|  | MLP | activation | relu |
|  |  | alpha | 0.05 |
|  |  | hidden_layer_sizes | (50, 50, 50) |
|  |  | learning_rate | constant |
|  |  | solver | adam |
|  | ET | bootstrap | False |
|  |  | criterion | gini |
|  |  | max_depth | 20 |
|  |  | max_features | sqrt |
|  |  | n_estimators | 200 |
|  | PLS | n_components | 2 |
|  |  | scale | True |
|  |  | max_itr | 500 |
|  |  | tol | 1e-06 |
| Meta layer | SVM | C | 1 |
|  |  | degree | 3 |
|  |  | gamma | scale |
|  |  | kernel | rbf |

### 3.7. Stacking ensemble significantly improves prediction performance

The best 5 feature combinations from Supplementary Table S1, with respect to F1-score, are highlighted in Table 7. In addition, FC-6 is the augmentation of FC-1 with the predictions obtained from the base layer predictors. In our stacking ensemble model, FC-6 is what the SVM learner in the meta layer learns on.

**Table 7:** Top 5 feature combinations from Table S1 in terms of F1-score. These combinations have been named from FC-1 to FC-5 for ease of reference in subsequent experiments. FC-6 is the augmentation of FC-1 with the base learner prediction values.

| Name | Feature Combination |
| --- | --- |
| FC-1 | ProtT5-XL-U50, PSSM |
| FC-2 | ProtT5-XL-U50, PSSM, TA, MG, DPC, ASA |
| FC-3 | ProtT5-XL-U50, PSSM, TA, MG, DPC |
| FC-4 | ProtT5-XL-U50, PSSM, TA |
| FC-5 | ProtT5-XL-U50, PSSM, TA, MG, DPC, PHYSICO |
| FC-6 | ProtT5-XL-U50, PSSM, BLP |

To examine the efficacy of stacking, we checked the performance of six SVM models trained using the validation set. Each model was trained using one of the six feature combinations mentioned in Table 7. The 10-fold CV performance comparison is shown in Figure 3. The FC-6 feature combination, which represents stacking, clearly outperforms the other combinations with respect to most of the performance metrics. Additionally, we used the t-distributed stochastic neighbor embedding (t-SNE) [58] to compare the feature space distribution of the feature combinations of Table 7 on the balanced validation dataset. As can be observed from Figure 4, the feature space of FC-6 produces a better separation between red and blue spots. This explains the superior predictive ability of the stacking ensemble compared to the other baseline models.

**Figure 3.**
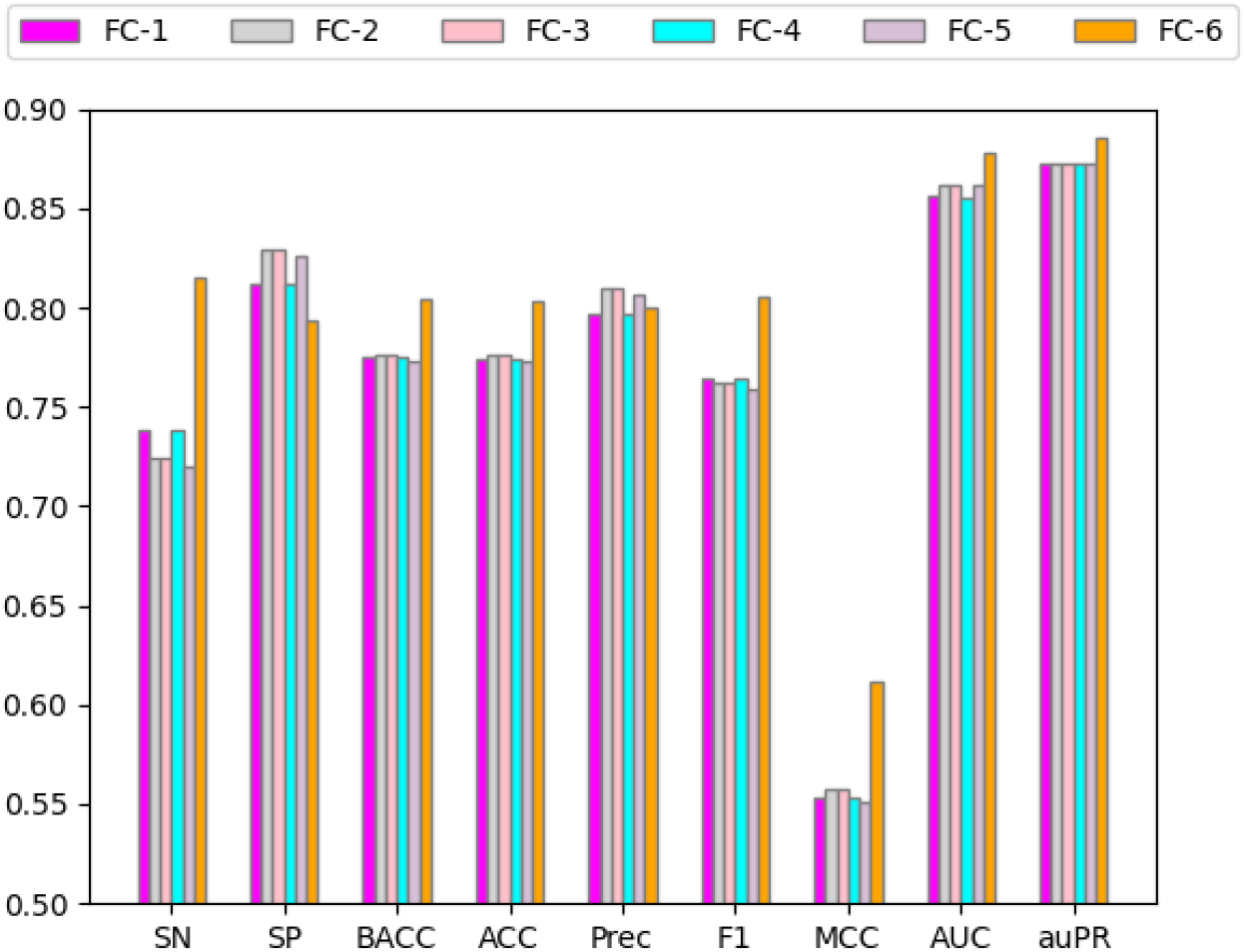
10-fold CV performance of SVM models trained on the validation set for various feature combinations. The validation set was balanced using random undersampling prior to training.

**Figure 4.**
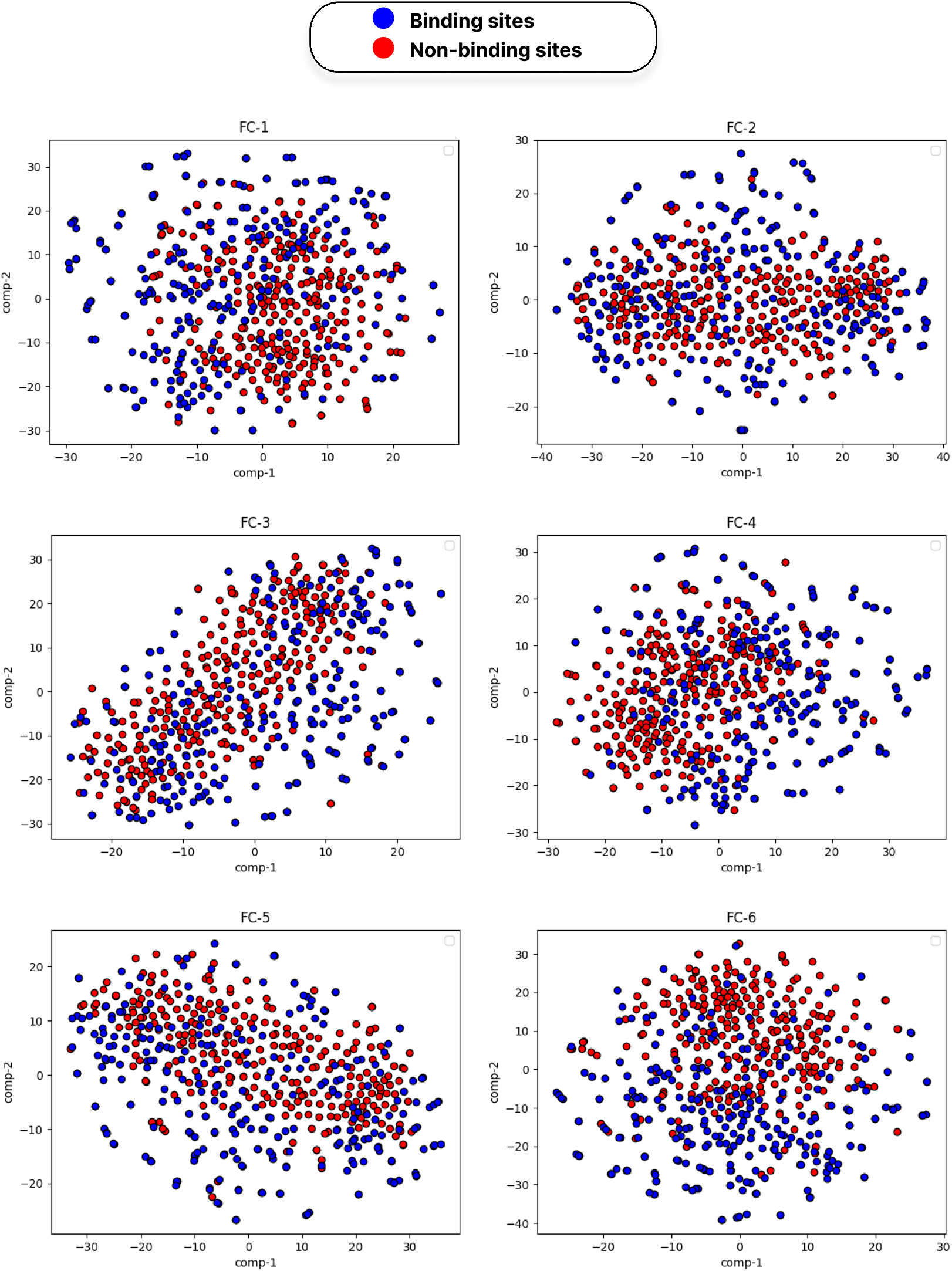
Six t-SNE plots on the validation dataset reflect the distribution of binding sites (blue dots) and non-binding sites (red dots) for the six feature combinations mentioned in Table 7. The feature space corresponding to FC-6 produces a much clearer separation between the binding and non-binding sites than other feature combinations.

### 3.8. StackCBEmbed outperforms state-of-the-art methods

We have used two independent test sets from the literature, TS49 and TS88, to compare the performance of StackCBEmbed with state-of-the-art (SOTA) methods, such as SPRINT-CBH, StackCBPred, PS-G, PS-S, CAPSIF:G, and CAPSIF:V. The SPRINT-CBH and StackCBPred servers were inaccessible, preventing us from reporting some of the metrics. Nevertheless, the trained model of StackCBPred was available in its GitHub repository, enabling us to replicate the results. We have presented both the published values as well as the reproduced values of performance metrics in Tables 8 and 9. For PS-G, PS-S, CAPSIF:G, and CAPSIF:V, we used the scripts provided in their respective GitHub repositories and produced their results on the TS49 and TS88 independent test sets.

**Table 8:** Comparison of StackCBEmbed with SOTA methods on the TS49 test set.

| Predictor | SN | SP | BACC | ACC | F1 | MCC | AUC | auPR |
| --- | --- | --- | --- | --- | --- | --- | --- | --- |
| StackCBPred (as per [20]) | 0.553 | 0.795 | 0.674 | 0.786 | - | 0.159 | - | - |
| StackCBPred (reproduced) | 0.522 | 0.803 | 0.662 | 0.792 | 0.157 | 0.151 | 0.725 | 0.106 |
| SPRINT-CBH | 0.389 | 0.925 | 0.657 | 0.906 | - | 0.195 | 0.755 | - |
| PS-G | 0.581 | 0.929 | 0.755 | 0.916 | 0.338 | 0.336 | **0.848** | **0.352** |
| PS-S | 0.274 | **0.986** | 0.63 | **0.96** | 0.335 | 0.325 | 0.809 | 0.273 |
| CAPSIF:G | 0.325 | 0.982 | 0.653 | 0.958 | 0.361 | 0.342 | 0.794 | 0.25 |
| CAPSIF:V | 0.429 | 0.975 | 0.702 | 0.954 | **0.411** | **0.387** | 0.753 | 0.288 |
| StackCBEmbed (PSSM) | 0.553 | 0.742 | 0.647 | 0.735 | 0.134 | 0.125 | 0.706 | 0.085 |
| StackCBEmbed (Embedding) | **0.736** | 0.817 | **0.777** | 0.814 | 0.227 | 0.260 | 0.834 | 0.237 |
| StackCBEmbed | 0.730 | 0.821 | 0.776 | 0.818 | 0.229 | 0.261 | 0.832 | 0.233 |

**Table 9:** Comparison of StackCBEmbed with SOTA methods on the TS88 test set.

| Predictor | SN | SP | BACC | ACC | F1 | MCC | AUC | auPR |
| --- | --- | --- | --- | --- | --- | --- | --- | --- |
| StackCBPred (as per [20]) | 0.565 | 0.797 | 0.681 | - | - | 0.139 | - | - |
| StackCBPred (reproduced) | 0.499 | 0.750 | 0.624 | 0.744 | 0.090 | 0.090 | 0.671 | 0.055 |
| SPRINT-CBH | 0.130 | **0.997** | 0.564 | - | - | 0.257 | - | - |
| PS-G | 0.513 | 0.922 | 0.717 | 0.911 | 0.227 | 0.24 | **0.823** | **0.233** |
| PS-S | 0.259 | 0.986 | 0.622 | **0.967** | **0.287** | **0.272** | 0.769 | 0.18 |
| CAPSIF:G | 0.267 | 0.973 | 0.62 | 0.955 | 0.233 | 0.212 | 0.715 | 0.111 |
| CAPSIF:V | 0.339 | 0.966 | 0.653 | 0.95 | 0.257 | 0.241 | 0.687 | 0.13 |
| StackCBEmbed (PSSM) | 0.494 | 0.740 | 0.617 | 0.733 | 0.086 | 0.083 | 0.668 | 0.047 |
| StackCBEmbed (Embedding) | 0.657 | 0.818 | 0.738 | 0.814 | 0.152 | 0.189 | 0.784 | 0.136 |
| StackCBEmbed | **0.666** | 0.818 | **0.742** | 0.814 | 0.154 | 0.192 | 0.788 | 0.141 |

While StackCBEmbed uses only two feature descriptors, PSSM and embedding, PSSM is expensive to compute. Therefore, we created two other models, one using PSSM only, and the other one only using the embedding. We wanted to check how well the model performs in case the PSSM for a query protein cannot be provided. From the results in Tables 8 and 9, we can see that when only PSSM is used, the model produces poor results in both datasets. However, when it uses embedding only, the results are quite good. In both datasets, it beats SOTA methods in terms of sensitivity and balanced accuracy. When both feature descriptors are combined, the results are further improved.

With SN scores of 73.6% and 66.6% and SP scores of 82.1% and 81.8%, StackCBEmbed achieved the highest BACC scores of 77.6% and 74.2% on TS49 and TS88 sets, respectively. It represents the model’s robust and consistent performance across both positive and negative class samples. This balance is significant in highly imbalanced datasets, and this makes StackCBEmbed a reliable and generalizable predictor. The model delivers AUC scores of 0.832 and 0.788 on TS49 and TS88 sets, respectively, which underlines its strong discriminating capability, even outperforming most predictors in this metric. StackCBEmbed is the most suitable predictor as it shows a balanced and consistent performance across both classes rather than just over-optimizing one class and neglecting the other one.

The Receiver Characteristic curve (ROC) represents the true positive rate (TPR) and false positive rate (FPR) values at several classification thresholds. Figure 5 shows the ROC curves (in independent test sets) for different flavors of StackCBEmbed. It also plots the same for other SOTA methods. From the curves in both datasets, it is clear that PSSM alone cannot give enough predictive capability to StackCBEmbed. However, when the embedding feature is added, its performance improves drastically. When embedding alone is used, the performance is very close to the regular StackCBEmbed. From the plots, we can see that StackCBEmbed beat other SOTA methods except PS-G. However, the performance of PS-G might be an overestimation for the reason described below.

**Figure 5.**
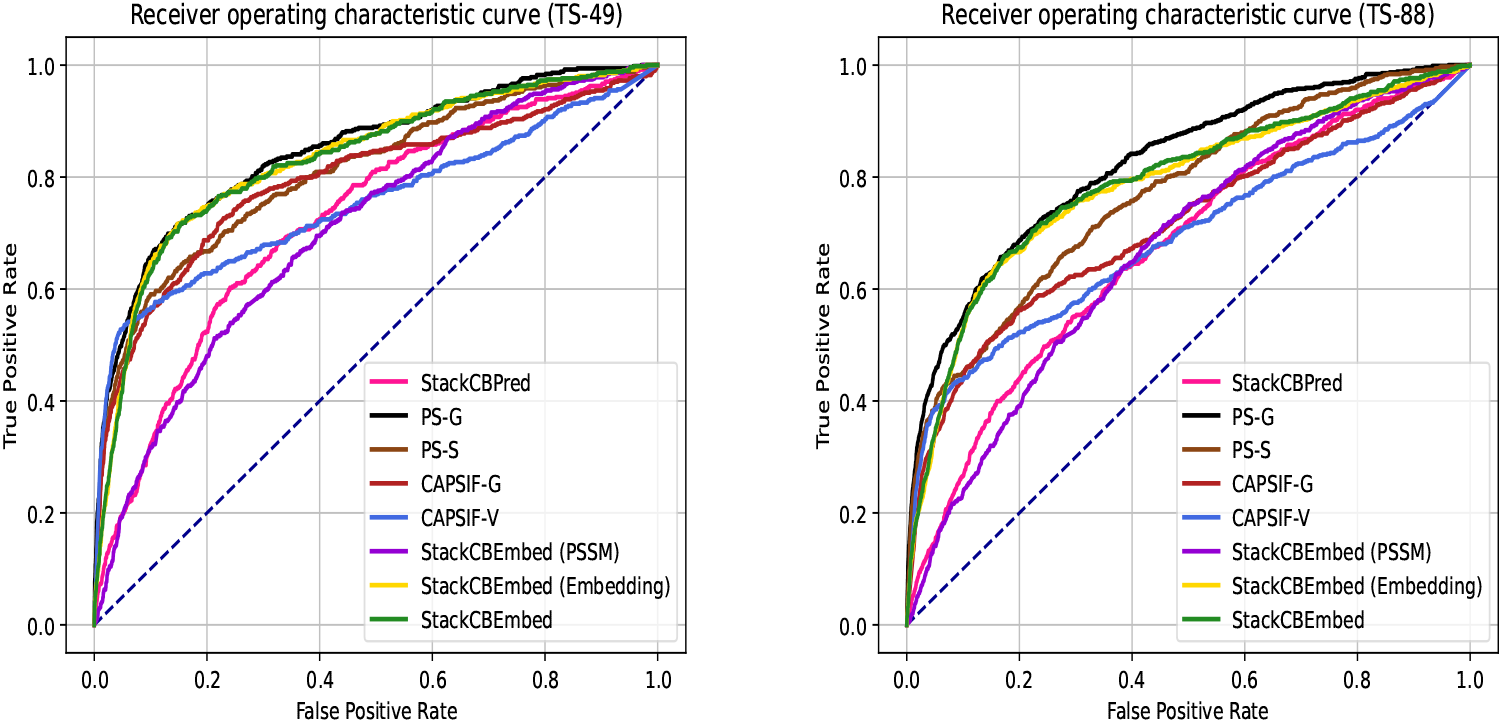
Comparison of Receiver Operating Characteristic (ROC) curves on TS49 and TS88 test sets.

The performance results of CAPSIF:G and CAPSIF:V predictors indicate significant overestimation due to substantial redundancy between the test sets used here and the training sets of these models. With 30% sequence similarity cutoff using BLAST-CLUST, we found that there were 18 redundant protein sequences in the TS49 set and 27 redundant protein sequences in the TS88 set for CAPSIF:G and CAPSIF:V. CAPSIF:V obtained SN scores of 38.5% and 29.0%, BACC scores of 67.9% and 62.6%, AUC scores of 71.5% and 65.1%, and auPR scores of 24.9% and 10.3% in non-redundant TS49 and TS88 sets (after removing the redundant protein sequences), respectively. CAPSIF:G achieved SN scores of 24.4% and 18.7%, BACC scores of 61.4% and 57.9%, AUC scores of 75.4% and 66.2%, and auPR scores of 21.1% and 7.4% in non-redundant TS49 and TS88 sets (removing the redundant protein sequences), respectively. These results highlight how crucial it is to make sure that the training and test sets do not have any redundancy to avoid overestimation. However, the training datasets for PS-G, and PS-S models were not properly provided in the respective GitHub repositories. We could not sort this out by contacting the authors either. As such, we could not measure the redundancy between their training sets, and TS88 and TS49; and the results for these models in Tables 8 and 9 are likely overestimations.

As both the test sets are heavily imbalanced, we also produced performance results after balancing the datasets. We kept all the positive samples and randomly undersampled the negative ones to make the dataset balanced. We did it 20 times and reported our results in mean ± stdev format in Tables 10 and 11. Not surprisingly, StackCBEmbed outperformed SOTA methods in SN, BACC, F1, and MCC metrics.

**Table 10:** Comparison of StackCBEmbed with SOTA methods on the TS49 test set balanced with random undersam-pling. Values are represented as mean ± stdev. Metrics are averaged over 20 replicates.

| Predictor | SN | SP | BACC | ACC | F1 | MCC | AUC | auPR |
| --- | --- | --- | --- | --- | --- | --- | --- | --- |
| StackCBPred (reproduced) | 0.522 ± 0.000 | 0.801 ± 0.017 | 0.661 ± 0.008 | 0.661 ± 0.008 | 0.606 ± 0.006 | 0.336 ± 0.019 | 0.724 ± 0.011 | 0.715 ± 0.017 |
| PS-G | 0.581 ± 0.000 | 0.926 ± 0.010 | 0.753 ± 0.005 | 0.753 ± 0.005 | 0.702 ± 0.004 | 0.540 ± 0.012 | **0.846** ± 0.008 | **0.862** ± 0.009 |
| PS-S | 0.274 ± 0.000 | **0.986** ± 0.005 | 0.630 ± 0.002 | 0.630 ± 0.002 | 0.425 ± 0.001 | 0.370 ± 0.009 | 0.806 ± 0.006 | 0.828 ± 0.009 |
| CAPSIF:G | 0.325 ± 0.000 | 0.980 ± 0.003 | 0.653 ± 0.002 | 0.653 ± 0.002 | 0.483 ± 0.001 | 0.404 ± 0.006 | 0.793 ± 0.005 | 0.820 ± 0.006 |
| CAPSIF:V | 0.429 ± 0.000 | 0.972 ± 0.006 | 0.701 ± 0.003 | 0.701 ± 0.003 | 0.589 ± 0.002 | 0.478 ± 0.009 | 0.751 ± 0.007 | 0.805 ± 0.008 |
| StackCBEmbed (PSSM) | 0.553 ± 0.000 | 0.739 ± 0.017 | 0.646 ± 0.008 | 0.646 ± 0.008 | 0.610 ± 0.006 | 0.298 ± 0.018 | 0.707 ± 0.010 | 0.688 ± 0.017 |
| StackCBEmbed (Embedding) | **0.736** ± 0.000 | 0.817 ± 0.018 | **0.777** ± 0.009 | **0.777** ± 0.009 | **0.767** ± 0.007 | **0.555** ± 0.019 | 0.833 ± 0.007 | 0.835 ± 0.012 |
| StackCBEmbed | 0.730 ± 0.000 | 0.820 ± 0.019 | 0.775 ± 0.010 | 0.775 ± 0.010 | 0.765 ± 0.008 | 0.552 ± 0.020 | 0.831 ± 0.007 | 0.833 ± 0.012 |

**Table 11:** Comparison of StackCBEmbed with SOTA methods on the random undersampled TS88 test set balanced with random undersampling. Values are represented as mean ± stdev. Metrics are averaged over 20 replicates.

| Predictor | SN | SP | BACC | ACC | F1 | MCC | AUC | auPR |
| --- | --- | --- | --- | --- | --- | --- | --- | --- |
| StackCBPred (reproduced) | 0.499 ± 0.000 | 0.755 ± 0.010 | 0.627 ± 0.005 | 0.627 ± 0.005 | 0.572 ± 0.003 | 0.262 ± 0.011 | 0.673 ± 0.008 | 0.666 ± 0.011 |
| PS-G | 0.513 ± 0.000 | 0.920 ± 0.011 | 0.717 ± 0.006 | 0.717 ± 0.006 | 0.644 ± 0.005 | 0.475 ± 0.015 | **0.822** ± 0.006 | **0.836** ± 0.009 |
| PS-S | 0.259 ± 0.000 | **0.985** ± 0.004 | 0.622 ± 0.002 | 0.622 ± 0.002 | 0.406 ± 0.001 | 0.354 ± 0.007 | 0.767 ± 0.007 | 0.788 ± 0.007 |
| CAPSIF:G | 0.267 ± 0.000 | 0.973 ± 0.007 | 0.620 ± 0.003 | 0.620 ± 0.003 | 0.413 ± 0.002 | 0.340 ± 0.013 | 0.715 ± 0.008 | 0.747 ± 0.011 |
| CAPSIF:V | 0.339 ± 0.000 | 0.966 ± 0.008 | 0.652 ± 0.004 | 0.652 ± 0.004 | 0.493 ± 0.003 | 0.391 ± 0.014 | 0.686 ± 0.005 | 0.741 ± 0.009 |
| StackCBEmbed (PSSM) | 0.494 ± 0.000 | 0.741 ± 0.012 | 0.618 ± 0.006 | 0.618 ± 0.006 | 0.564 ± 0.004 | 0.243 ± 0.013 | 0.670 ± 0.008 | 0.644 ± 0.012 |
| StackCBEmbed (Embedding) | 0.657 ± 0.000 | 0.816 ± 0.012 | 0.736 ± 0.006 | 0.736 ± 0.006 | 0.714 ± 0.005 | 0.479 ± 0.013 | 0.782 ± 0.007 | 0.791 ± 0.012 |
| StackCBEmbed | **0.666** ± 0.000 | 0.814 ± 0.010 | **0.740** ± 0.005 | **0.740** ± 0.005 | **0.719** ± 0.004 | **0.485** ± 0.011 | 0.786 ± 0.006 | 0.795 ± 0.012 |

### 3.9. StackCBEmbed performed well on TS46 test set

We have tested the different variations of StackCBEmbed on a third independent test set, TS46. The performance results are shown in Tables 12 and 13. It performed significantly well on this test set as well. This is a further testament to the generalizability of StackCBEmbed in unseen data.

**Table 12:**
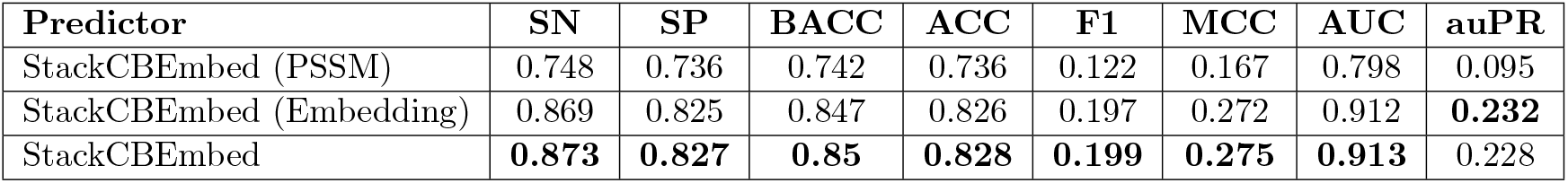
Performance of StackCBEmbed on the TS46 test set.

| Predictor | SN | SP | BACC | ACC | F1 | MCC | AUC | auPR |
| --- | --- | --- | --- | --- | --- | --- | --- | --- |
| StackCBEmbed (PSSM) | 0.748 | 0.736 | 0.742 | 0.736 | 0.122 | 0.167 | 0.798 | 0.095 |
| StackCBEmbed (Embedding) | 0.869 | 0.825 | 0.847 | 0.826 | 0.197 | 0.272 | 0.912 | **0.232** |
| StackCBEmbed | **0.873** | **0.827** | **0.85** | **0.828** | **0.199** | **0.275** | **0.913** | 0.228 |

**Table 13:**
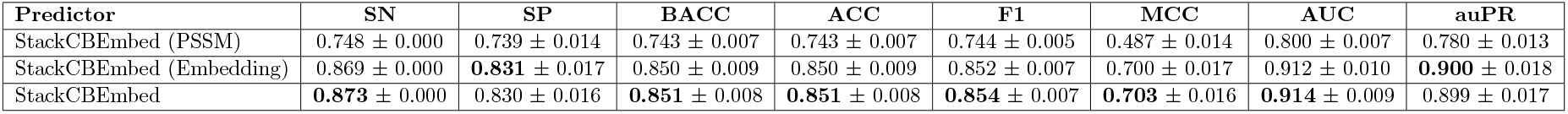
Performance of StackCBEmbed on the TS46 test set balanced with random undersampling. Values are represented as mean ± stdev. Metrics are averaged over 20 replicates.

| Predictor | SN | SP | BACC | ACC | F1 | MCC | AUC | auPR |
| --- | --- | --- | --- | --- | --- | --- | --- | --- |
| StackCBEmbed (PSSM) | 0.748 ± 0.000 | 0.739 ± 0.014 | 0.743 ± 0.007 | 0.743 ± 0.007 | 0.744 ± 0.005 | 0.487 ± 0.014 | 0.800 ± 0.007 | 0.780 ± 0.013 |
| StackCBEmbed (Embedding) | 0.869 ± 0.000 | **0.831** ± 0.017 | 0.850 ± 0.009 | 0.850 ± 0.009 | 0.852 ± 0.007 | 0.700 ± 0.017 | 0.912 ± 0.010 | **0.900** ± 0.018 |
| StackCBEmbed | **0.873** ± 0.000 | 0.830 ± 0.016 | **0.851** ± 0.008 | **0.851** ± 0.008 | **0.854** ± 0.007 | **0.703** ± 0.016 | **0.914** ± 0.009 | 0.899 ± 0.017 |

## 4. Discussion and Conclusion

In this study, we have proposed a novel prediction model for protein-carbohydrate binding sites. Named StackCBEmbed, the predictor is a stacking ensemble model trained using 98 protein sequences (≤ 30% pairwise sequence identity), comprising 1014 binding and 25705 non-binding residues. 70% of this data (training set) was used to train the base layer, while the remaining 30% (validation set) was used to train the meta layer. From a variety of traditional sequence-based features, and embedding from recently trained protein language models, the optimal feature vector was prepared using incremental feature selection. In the final model, a combination of PSSM (20 features) and embedding features (1024 features) were used. Therefore each residue was represented as a 1024 + 20 = 1044 dimensional feature vector. As the training data is heavily imbalanced, for improved learning, 10 different subsamples was prepared, each with a different ratio of positive and negative samples. The base layer of the model used SVM, MLP, ET, and PLS learners. As the meta layer, an SVM classifier was trained on the validation set, augmented with the prediction probabilities from the base layer models, and balanced using random undersampling. Results in two independent test sets indicate that StackCBEmbed outperforms SOTA methods in terms of BACC and SN metrics even when only the embedding features are used. The results further improve if PSSM features are added. This is an important observation because PSSM cannot be computed for proteins with not enough homologs in the database. Also, for proteins with enough homologs, the process to compute PSSM is time-consuming. Therefore, a researcher can choose to avoid using PSSM and still get good prediction results using StackCBEmbed.

We also looked at another SOTA method stemming out of the work by Malik et al. [6]. This publication is quite old and uses a dataset of 40 protein-carbohydrate complexes only. Neither a prediction server nor a standalone downloadable version was available for this method. Therefore we opted to exclude this approach from our benchmarking. More recent methods, such as, LMNGlypred [59] and PUStackNGly [60] were also reviewed. But these models are primarily designed for predicting N-linked glycosylation sites. Our work, on the other hand, focuses on identifying non-specific carbohydrate binding sites, rendering these methods out of scope for comparative analysis.

Prediction models in different problems in bioinformatics are dominated by deep learning methods in the modern era. Accordingly, we started our exploration with several deep learning architectures alongside the traditional ML algorithms. Specifically, we investigated CNN, RNN, LSTM and Transformer models that are well-suited for sequence data. However, these did not yield promising results in this prediction problem (See Supplementary Tables S2 and S3). This might have been due to the small size of the dataset and significant imbalance therein. In fact, many recent studies [61, 62, 63, 64, 65] have used traditional machine learning models to surpass the performance of deep learning (DL) based methods. DL models do not perform well under small datasets as they tend to have millions (or even billions) of parameters to learn, resulting in overfitting [66]. DL models are expensive during inference as well. Therefore, the contribution of traditional ML in the field of bioinformatics is still very pertinent.

In our study, we explored and analyzed sequence-derived structural features. We could not generate predicted PDBs from AlphaFold [67] due to its high computational demands and resource-intensive nature. For instance, generating a predicted PDB file for the protein sequence ID 1F31A, having a sequence length of 1280 residues, took 3.72 hours with 16 GB of memory and 15 GB of VRAM. Although this limited our ability to incorporate AlphaFold-generated predicted PDBs in our model, we acknowledge the potential of using structural features from predicted PDBs to predict protein-carbohydrate binding sites and leave it as future work.

While StackCBEmbed outperformed SOTA methods in BACC and SN metrics in both the independent test sets, analysis of F1 and auPR scores revealed room for improvement. In particular, the tool needs to improve its precision. These findings highlight the importance of further research and development in this domain. StackCBEmbed achieved a balanced accuracy and sensitivity of 77.6% and 73%, respectively, in the TS49 test set. In the case of TS88, these values were 74.2% and 66.6%, respectively. Thus StackCBEmbed brings in significant improvements in protein-carbohydrate binding site prediction. The model is freely available as an easy to use python script at https://github.com/nafiislam/StackCBEmbed. We hope that research related to protein-carbohydrate binding sites will immensely benefit from the efficacy of this novel, easy-to-use predictor.

## Declarations

### Ethics approval and consent to participate

Not applicable

### Consent for publication

Not applicable

### Availability of data and materials

The StackCBEmbed model, associated data and scripts to reproduce the experimental results are available publicly at https://github.com/nafiislam/StackCBEmbed.

### Competing interests

The authors declare that they have no competing interests

### Funding

MSR is partially supported by Basic Research Grant administered by BUET. MMIN is supported by the RISE Student Research Grant [No. S2024-01-004], administered by RISE, BUET.

### Authors’ contributions

MSR and QFN conceived the research. TNI did preliminary explorations. MSR, QFN, and MMIN set up the experiments. MMIN and QFN conducted rigorous explorations to finalize the prediction model. MMIN validated all the experiments. MSR, QFN, and MMIN analyzed all the results. MMIN improved the code and wrote scripts to reproduce each experiment result. All authors reviewed and approved the manuscript.

## Acknowledgements

Not applicable

## Supplementary Material

This supplementary section provides additional tables.

**Table S1:** 10-fold CV performance of XGB predictor trained on the benchmark dataset (balanced with random undersampling) using various feature subsets as per Algorithm 1. An extra horizontal line separates each increment phase. ✓mark indicates that the row was selected for the next increment phase (based on F1-score). The best F1-score across all subsets is bold-faced.

| Feature combination | SN | SP | BACC | Acc | Prec | F1 | MCC | AUC | auPR |
| --- | --- | --- | --- | --- | --- | --- | --- | --- | --- |
| PSSM | 0.641 | 0.677 | 0.659 | 0.659 | 0.666 | 0.653 | 0.318 | 0.726 | 0.726 |
| MonoGram | 0.576 | 0.550 | 0.563 | 0.563 | 0.562 | 0.568 | 0.127 | 0.603 | 0.613 |
| DPC | 0.670 | 0.685 | 0.678 | 0.678 | 0.681 | 0.675 | 0.356 | 0.758 | 0.761 |
| ASA | 0.561 | 0.468 | 0.515 | 0.515 | 0.515 | 0.536 | 0.030 | 0.521 | 0.516 |
| HSE | 0.605 | 0.549 | 0.577 | 0.577 | 0.573 | 0.588 | 0.154 | 0.608 | 0.595 |
| Torsion_angles | 0.564 | 0.516 | 0.540 | 0.540 | 0.540 | 0.551 | 0.080 | 0.568 | 0.554 |
| Physicochemical | 0.492 | 0.535 | 0.514 | 0.514 | 0.514 | 0.501 | 0.027 | 0.516 | 0.521 |
| ProtT5-XL-U50 | 0.754 | 0.810 | 0.782 | 0.782 | 0.800 | 0.776 | 0.566 | 0.864 | 0.875 |
| ProtT5-XL-U50, PSSM | 0.776 | 0.814 | 0.795 | 0.795 | 0.806 | **0.791** | 0.591 | 0.869 | 0.879 |
| ProtT5-XL-U50, MonoGram | 0.759 | 0.804 | 0.782 | 0.782 | 0.795 | 0.776 | 0.564 | 0.868 | 0.880 |
| ProtT5-XL-U50, DPC | 0.772 | 0.801 | 0.786 | 0.787 | 0.795 | 0.783 | 0.574 | 0.869 | 0.878 |
| ProtT5-XL-U50, ASA | 0.758 | 0.815 | 0.787 | 0.787 | 0.806 | 0.781 | 0.576 | 0.869 | 0.877 |
| ProtT5-XL-U50, HSE | 0.773 | 0.801 | 0.787 | 0.787 | 0.795 | 0.784 | 0.575 | 0.865 | 0.875 |
| ProtT5-XL-U50, Torsion_angles | 0.766 | 0.816 | 0.791 | 0.791 | 0.806 | 0.785 | 0.583 | 0.865 | 0.874 |
| ProtT5-XL-U50, Physicochemical | 0.766 | 0.812 | 0.789 | 0.789 | 0.804 | 0.784 | 0.579 | 0.866 | 0.877 |
| ProtT5-XL-U50, PSSM, MonoGram | 0.769 | 0.815 | 0.792 | 0.792 | 0.808 | 0.787 | 0.586 | 0.867 | 0.879 |
| ProtT5-XL-U50, PSSM, DPC | 0.767 | 0.817 | 0.792 | 0.792 | 0.810 | 0.787 | 0.587 | 0.870 | 0.882 |
| ProtT5-XL-U50, PSSM, ASA | 0.764 | 0.807 | 0.785 | 0.786 | 0.799 | 0.781 | 0.572 | 0.869 | 0.879 |
| ProtT5-XL-U50, PSSM, HSE | 0.778 | 0.804 | 0.791 | 0.791 | 0.799 | 0.788 | 0.582 | 0.864 | 0.876 |
| ProtT5-XL-U50, PSSM, Torsion_angles | 0.771 | 0.816 | 0.793 | 0.793 | 0.808 | 0.788 | 0.588 | 0.870 | 0.878 |
| ProtT5-XL-U50, PSSM, Physicochemical | 0.763 | 0.819 | 0.791 | 0.791 | 0.808 | 0.785 | 0.583 | 0.862 | 0.875 |
| ProtT5-XL-U50, PSSM, Torsion_angles, MonoGram | 0.770 | 0.814 | 0.792 | 0.792 | 0.807 | 0.787 | 0.585 | 0.868 | 0.879 |
| ProtT5-XL-U50, PSSM, Torsion_angles, DPC | 0.762 | 0.803 | 0.782 | 0.783 | 0.795 | 0.778 | 0.566 | 0.870 | 0.879 |
| ProtT5-XL-U50, PSSM, Torsion_angles, ASA | 0.768 | 0.805 | 0.786 | 0.787 | 0.798 | 0.782 | 0.574 | 0.864 | 0.874 |
| ProtT5-XL-U50, PSSM, Torsion_angles, HSE | 0.761 | 0.802 | 0.782 | 0.782 | 0.794 | 0.777 | 0.564 | 0.861 | 0.878 |
| ProtT5-XL-U50, PSSM, Torsion_angles, Physicochemical | 0.765 | 0.818 | 0.792 | 0.792 | 0.809 | 0.786 | 0.585 | 0.863 | 0.871 |
| ProtT5-XL-U50, PSSM, Torsion_angles, MonoGram, DPC | 0.771 | 0.818 | 0.795 | 0.795 | 0.810 | 0.789 | 0.591 | 0.871 | 0.878 |
| ProtT5-XL-U50, PSSM, Torsion_angles, MonoGram, ASA | 0.765 | 0.807 | 0.786 | 0.786 | 0.798 | 0.781 | 0.573 | 0.864 | 0.874 |
| ProtT5-XL-U50, PSSM, Torsion_angles, MonoGram, HSE | 0.770 | 0.804 | 0.787 | 0.787 | 0.797 | 0.783 | 0.575 | 0.871 | 0.881 |
| ProtT5-XL-U50, PSSM, Torsion_angles, MonoGram, Physicochemical | 0.769 | 0.806 | 0.787 | 0.788 | 0.799 | 0.783 | 0.576 | 0.862 | 0.874 |
| ProtT5-XL-U50, PSSM, Torsion_angles, MonoGram, DPC, ASA | 0.769 | 0.822 | 0.796 | 0.796 | 0.813 | 0.790 | 0.593 | 0.872 | 0.877 |
| ProtT5-XL-U50, PSSM, Torsion_angles, MonoGram, DPC, HSE | 0.773 | 0.811 | 0.792 | 0.792 | 0.803 | 0.787 | 0.585 | 0.870 | 0.871 |
| ProtT5-XL-U50, PSSM, Torsion_angles, MonoGram, DPC, Physicochemical | 0.773 | 0.812 | 0.792 | 0.792 | 0.805 | 0.788 | 0.586 | 0.868 | 0.876 |
| ProtT5-XL-U50, PSSM, Torsion_angles, MonoGram, DPC, ASA, HSE | 0.755 | 0.814 | 0.784 | 0.785 | 0.803 | 0.778 | 0.571 | 0.871 | 0.877 |
| ProtT5-XL-U50, PSSM, Torsion_angles, MonoGram, DPC, ASA, Physicochemical | 0.769 | 0.817 | 0.793 | 0.793 | 0.809 | 0.788 | 0.588 | 0.870 | 0.873 |
| ProtT5-XL-U50, PSSM, Torsion_angles, MonoGram, DPC, ASA, Physicochemical, HSE | 0.765 | 0.819 | 0.792 | 0.792 | 0.808 | 0.786 | 0.585 | 0.874 | 0.879 |

**Table S2:** Comparison of several deep learning based models with StackCBEmbed on the TS49 dataset.

| **Predictor** | **SN** | **SP** | **BACC** | **ACC** | **F1** | **MCC** | **AUC** | **auPR** |
| --- | --- | --- | --- | --- | --- | --- | --- | --- |
| StackCBEmbed | **0.73** | **0.821** | **0.776** | **0.818** | **0.229** | **0.261** | **0.832** | **0.233** |
| CNN | 0.341 | 0.736 | 0.538 | 0.721 | 0.083 | 0.032 | 0.584 | 0.046 |
| RNN | 0.307 | 0.739 | 0.523 | 0.723 | 0.076 | 0.02 | 0.541 | 0.041 |
| LSTM | 0.394 | 0.634 | 0.514 | 0.626 | 0.072 | 0.011 | 0.534 | 0.041 |
| Transformer | 0.384 | 0.682 | 0.533 | 0.671 | 0.079 | 0.026 | 0.556 | 0.046 |

**Table S3:** Comparison of several deep learning based models with StackCBEmbed on the TS88 set.

| **Predictor** | **SN** | **SP** | **BACC** | **ACC** | **F1** | **MCC** | **AUC** | **auPR** |
| --- | --- | --- | --- | --- | --- | --- | --- | --- |
| StackCBEmbed | **0.666** | **0.818** | **0.742** | **0.814** | **0.154** | **0.192** | **0.788** | **0.141** |
| CNN | 0.345 | 0.796 | 0.571 | 0.785 | 0.075 | 0.055 | 0.607 | 0.053 |
| RNN | 0.275 | 0.744 | 0.509 | 0.732 | 0.049 | 0.007 | 0.541 | 0.028 |
| LSTM | 0.388 | 0.641 | 0.515 | 0.635 | 0.051 | 0.010 | 0.540 | 0.027 |
| Transformer | 0.448 | 0.603 | 0.525 | 0.599 | 0.054 | 0.016 | 0.559 | 0.036 |

## Notes

### Competing Interest Statement

The authors have declared no competing interest.

### Summary of Updates

First, we have improved the manuscript's organization and readability. The large Table 4, which disrupted the flow of the main text, has been moved to the Supplementary Material section. This allows readers to focus on the key methodological and result-oriented discussions in the main body while still having access to detailed information. Second, we have strengthened the justification for our choice of protein language model embeddings. A more comprehensive comparative analysis of different embeddings has been included in Section 3.1. In addition to the previously evaluated models, we have now incorporated ESM2 into the benchmarking. The results confirm that ProtT5-XL-U50 remains the most effective embedding model for the protein carbohydrate binding site prediction task considered in this study. Third, we have expanded the experimental evaluation by adding additional baseline models. To address concerns regarding residue-level contextual modeling, we implemented and evaluated CNN, RNN, LSTM, and Transformer-based architectures using protein language model embeddings. These results are now reported in the Supplementary Material, and their limitations are discussed in the Discussion section, particularly in relation to dataset size and class imbalance. Fourth, we have significantly extended the comparison with state-of-the-art methods. In addition to SPRINT-CBH and StackCBPred, we now include comparisons with several recently proposed approaches, namely PS-G, PS-S, CAPSIF:G, and CAPSIF:V. Section 3.8 has been updated accordingly to provide a more comprehensive and up-to-date performance evaluation. Fifth, we have clarified the novelty and contribution of this work by emphasizing the methodological innovation of combining stacking ensemble learning with protein language model embeddings for carbohydrate binding site prediction. We also highlight the preparation of a novel independent test dataset to assess generalizability, which will be useful for future benchmarking studies. Finally, we have expanded the Discussion section to address structure-based analysis. While structural information and AlphaFold-generated models can enhance interpretability, we explain why this work focuses on sequence-based prediction and outline structure-based extensions as future work. Overall, these revisions improve the clarity, robustness, and relevance of the manuscript while addressing all major reviewer concerns.

## References

[1] Clara Shionyu-Mitsuyama, Tsuyoshi Shirai, Hirokazu Ishida, and Takashi Yamane. An empirical approach for structure-based prediction of carbohydrate-binding sites on proteins. Protein Engineering, 16(7):467–478, 2003.

[2] Maria del Carmen Fernandez-Alonso, Dolores Díaz, Manuel Alvaro Berbis, Filipa Marcelo, Jesus Jimenez-Barbero, et al. Protein-carbohydrate interactions studied by NMR: from molecular recognition to drug design. Current Protein and Peptide Science, 13(8):816–830, 2012.

[3] Injae Shin, Sungjin Park, and Myung-ryul Lee. Carbohydrate microarrays: an advanced technology for functional studies of glycans. Chemistry–A European Journal, 11(10):2894–2901, 2005.

[4] Michaela Wimmerová, Stanislav Kozmon, Ivona Nečasová, Sushil Kumar Mishra, Jan Komárek, and Jaroslav Koča. Stacking interactions between carbohydrate and protein quantified by combination of theoretical and experimental methods. 2012.

[5] Ben Rathje, Caelen Begg, Liv Helland, and Pari Kyars. A review of common shoulder injuries: clavicular fractures and anterior dislocations. MacEwan University Student eJournal, 4(1), 2020.

[6] Adeel Malik and Shandar Ahmad. Sequence and structural features of carbohydrate binding in proteins and assessment of predictability using a neural network. BMC Structural Biology, 7:1–14, 2007.

[7] Alan Brown and Matthew K Higgins. Carbohydrate binding molecules in malaria pathology. Current opinion in structural biology, 20(5):560–566, 2010.

[8] KO François and Jan Balzarini. Potential of carbohydrate-binding agents as therapeutics against enveloped viruses. Medicinal research reviews, 32(2):349–387, 2012.

[9] Avraham Raz and Susumu Nakahara. Biological modulation by lectins and their ligands in tumor progression and metastasis. Anti-Cancer Agents in Medicinal Chemistry (Formerly Current Medicinal Chemistry-Anti-Cancer Agents), 8(1):22–36, 2008.

[10] Ghazaleh Taherzadeh, Yaoqi Zhou, Alan Wee-Chung Liew, and Yuedong Yang. Sequence-based prediction of protein–carbohydrate binding sites using support vector machines. Journal of chemical information and modeling, 56(10):2115–2122, 2016.

[11] Chiara Taroni, Susan Jones, and Janet M Thornton. Analysis and prediction of carbohydrate binding sites. Protein Engineering, 13(2):89–98, 2000.

[12] MS Sujatha and Petety V Balaji. Identification of common structural features of binding sites in galactose-specific proteins. Proteins: Structure, Function, and Bioinformatics, 55(1):44–65, 2004.

[13] Mahesh Kulharia, Stephen J Bridgett, Roger S Goody, and Richard M Jackson. InCa-SiteFinder: a method for structure-based prediction of inositol and carbohydrate binding sites on proteins. Journal of Molecular Graphics and Modelling, 28(3):297–303, 2009.

[14] Houssam Nassif, Hassan Al-Ali, Sawsan Khuri, and Walid Keirouz. Prediction of protein-glucose binding sites using support vector machines. Proteins: Structure, Function, and Bioinformatics, 77(1):121–132, 2009.

[15] Keng-Chang Tsai, Jhih-Wei Jian, Ei-Wen Yang, Po-Chiang Hsu, Hung-Pin Peng, Ching-Tai Chen, Jun-Bo Chen, Jeng-Yih Chang, Wen-Lian Hsu, and An-Suei Yang. Prediction of carbohydrate binding sites on protein surfaces with 3-dimensional probability density distributions of interacting atoms. PLoS One, 7(7):e40846, 2012.

[16] M Michael Gromiha, K Veluraja, and Kazuhiko Fukui. Identification and analysis of binding site residues in proteincarbohydrate complexes using energy based approach. Protein and Peptide Letters, 21(8):799–807, 2014.

[17] NR Siva Shanmugam, J Fermin Angelo Selvin, K Veluraja, and M Michael Gromiha. Identification and analysis of key residues involved in folding and binding of protein-carbohydrate complexes. Protein and peptide letters, 25(4):379–389, 2018.

[18] Priyadarshini P Pai and Sukanta Mondal. MOWGLI: prediction of protein–MannOse interacting residues With ensemble classifiers usinG evoLutionary Information. Journal of Biomolecular Structure and Dynamics, 34(10):2069–2083, 2016.

[19] Sandhya Agarwal, Nitish Kumar Mishra, Harinder Singh, and Gajendra PS Raghava. Identification of mannose interacting residues using local composition. PLoS One, 6(9):e24039, 2011.

[20] Suraj Gattani, Avdesh Mishra, and Md Tamjidul Hoque. StackCBPred: A stacking based prediction of protein-carbohydrate binding sites from sequence. Carbohydrate research, 486:107857, 2019.

[21] Samuel W Canner, Sudhanshu Shanker, and Jeffrey J Gray. Structure-based neural network protein–carbohydrate interaction predictions at the residue level. Frontiers in Bioinformatics, 3:1186531, 2023.

[22] Parth Bibekar, Lucien Krapp, and Matteo Dal Peraro. PeSTo-Carbs: Geometric Deep Learning for Prediction of Protein–Carbohydrate Binding Interfaces. Journal of Chemical Theory and Computation, 20(8):2985–2991, 2024.

[23] Tristan Bepler and Bonnie Berger. Learning the protein language: Evolution, structure, and function. Cell systems, 12(6):654–669, 2021.

[24] Ahmed Elnaggar, Michael Heinzinger, Christian Dallago, Ghalia Rehawi, Yu Wang, Llion Jones, Tom Gibbs, Tamas Feher, Christoph Angerer, Martin Steinegger, et al. ProtTrans: Toward Understanding the Language of Life Through Self-Supervised Learning. IEEE transactions on pattern analysis and machine intelligence, 44(10):7112–7127, 2021.

[25] Huiying Zhao, Yuedong Yang, Mark von Itzstein, and Yaoqi Zhou. Carbohydrate-binding protein identification by coupling structural similarity searching with binding affinity prediction. Journal of computational chemistry, 35(30):2177–2183, 2014.

[26] Adeel Malik, Ahmad Firoz, Vivekanand Jha, and Shandar Ahmad. PROCARB: a database of known and modelled carbohydrate-binding protein structures with sequence-based prediction tools. Advances in bioinformatics, 2010, 2010.

[27] Mark Johnson, Irena Zaretskaya, Yan Raytselis, Yuri Merezhuk, Scott McGinnis, and Thomas L Madden. NCBI BLAST: a better web interface. Nucleic acids research, 36(uppl 2):W5–W9, 2008.

[28] In-Kwon Yeo and Richard A Johnson. A new family of power transformations to improve normality or symmetry. Biometrika, 87(4):954–959, 2000.

[29] Sumaiya Iqbal and Md Tamjidul Hoque. PBRpredict-Suite: a suite of models to predict peptide-recognition domain residues from protein sequence. Bioinformatics, 34(19):3289–3299, 2018.

[30] Avdesh Mishra, Pujan Pokhrel, and Md Tamjidul Hoque. StackDPPred: a stacking based prediction of DNA-binding protein from sequence. Bioinformatics, 35(3):433–441, 2019.

[31] Ashis Kumer Biswas, Nasimul Noman, and Abdur Rahman Sikder. Machine learning approach to predict protein phosphorylation sites by incorporating evolutionary information. BMC bioinformatics, 11:1–17, 2010.

[32] Md Nasrul Islam, Sumaiya Iqbal, Ataur R Katebi, and Md Tamjidul Hoque. A balanced secondary structure predictor. Journal of theoretical biology, 389:60–71, 2016.

[33] Sumaiya Iqbal, Avdesh Mishra, and Md Tamjidul Hoque. Improved prediction of accessible surface area results in efficient energy function application. Journal of theoretical biology, 380:380–391, 2015.

[34] Ruchi Verma, Grish C Varshney, and GPS Raghava. Prediction of mitochondrial proteins of malaria parasite using split amino acid composition and PSSM profile. Amino acids, 39:101– 110, 2010.

[35] Stephen F Altschul, Warren Gish, Webb Miller, Eugene W Myers, and David J Lipman. Basic local alignment search tool. Journal of molecular biology, 215(3):403–410, 1990.

[36] Harsh Saini, Gaurav Raicar, Alok Sharma, Sunil Lal, Abdollah Dehzangi, Rajeshkannan Ananthanarayanan, James Lyons, Neela Biswas, and Kuldip K Paliwal. Protein structural class prediction via k-separated bigrams using position specific scoring matrix. Journal of Advanced Computational Intelligence and Intelligent Informatics, 18(4):474–479, 2014.

[37] Alok Sharma, James Lyons, Abdollah Dehzangi, and Kuldip K Paliwal. A feature extraction technique using bi-gram probabilities of position specific scoring matrix for protein fold recognition. Journal of theoretical biology, 320:41–46, 2013.

[38] Taigang Liu, Xiaoqi Zheng, and Jun Wang. Prediction of protein structural class for low-similarity sequences using support vector machine and PSI-BLAST profile. Biochimie, 92(10):1330–1334, 2010.

[39] Eshel Faraggi, Tuo Zhang, Yuedong Yang, Lukasz Kurgan, and Yaoqi Zhou. SPINE X: improving protein secondary structure prediction by multistep learning coupled with prediction of solvent accessible surface area and backbone torsion angles. Journal of computational chemistry, 33(3):259–267, 2012.

[40] Rhys Heffernan, Abdollah Dehzangi, James Lyons, Kuldip Paliwal, Alok Sharma, Jihua Wang, Abdul Sattar, Yaoqi Zhou, and Yuedong Yang. Highly accurate sequence-based prediction of half-sphere exposures of amino acid residues in proteins. Bioinformatics, 32(6):843–849, 2016.

[41] Rhys Heffernan, Kuldip Paliwal, James Lyons, Jaswinder Singh, Yuedong Yang, and Yaoqi Zhou. Single-sequence-based prediction of protein secondary structures and solvent accessibility by deep whole-sequence learning. Journal of computational chemistry, 39(26):2210–2216, 2018.

[42] Rhys Heffernan, Yuedong Yang, Kuldip Paliwal, and Yaoqi Zhou. Capturing non-local interactions by long short-term memory bidirectional recurrent neural networks for improving prediction of protein secondary structure, backbone angles, contact numbers and solvent accessibility. Bioinformatics, 33(18):2842–2849, 2017.

[43] Jiangning Song, Hao Tan, Kazuhiro Takemoto, and Tatsuya Akutsu. HSEpred: predict half-sphere exposure from protein sequences. Bioinformatics, 24(13):1489–1497, 2008.

[44] Jens Meiler, Michael Müller, Anita Zeidler, and Felix Schmäschke. Generation and evaluation of dimension-reduced amino acid parameter representations by artificial neural networks. Molecular modeling annual, 7(9):360–369, 2001.

[45] Zeming Lin, Halil Akin, Roshan Rao, Brian Hie, Zhongkai Zhu, Wenting Lu, Nikita Smetanin, Robert Verkuil, Ori Kabeli, Yaniv Shmueli, et al. Evolutionary-scale prediction of atomic-level protein structure with a language model. Science, 379(6637):1123–1130, 2023.

[46] Tom Brown, Benjamin Mann, Nick Ryder, Melanie Subbiah, Jared D Kaplan, Prafulla Dhariwal, Arvind Neelakantan, Pranav Shyam, Girish Sastry, Amanda Askell, et al. Language models are few-shot learners. Advances in neural information processing systems, 33:1877–1901, 2020.

[47] Zhenzhong Lan, Mingda Chen, Sebastian Goodman, Kevin Gimpel, Piyush Sharma, and Radu Soricut. Albert: A lite bert for self-supervised learning of language representations. arXiv preprint 1909.11942, 2019.

[48] Zhilin Yang, Zihang Dai, Yiming Yang, Jaime Carbonell, Russ R Salakhutdinov, and Quoc V Le. Xlnet: Generalized autoregressive pretraining for language understanding. Advances in neural information processing systems, 32, 2019.

[49] Colin Raffel, Noam Shazeer, Adam Roberts, Katherine Lee, Sharan Narang, Michael Matena, Yanqi Zhou, Wei Li, and Peter J Liu. Exploring the limits of transfer learning with a unified text-to-text transformer. The Journal of Machine Learning Research, 21(1):5485–5551, 2020.

[50] Douglas G Altman and J Martin Bland. Diagnostic tests. 1: Sensitivity and specificity. BMJ: British Medical Journal, 308(6943):1552, 1994.

[51] David MW Powers. Evaluation: from precision, recall and F-measure to ROC, informedness, markedness and correlation. arXiv preprint 2010.16061, 2020.

[52] Tom Fawcett. An introduction to ROC analysis. Pattern recognition letters, 27(8):861–874, 2006.

[53] Jens Keilwagen, Ivo Grosse, and Jan Grau. Area under precision-recall curves for weighted and unweighted data. PloS one, 9(3):e92209, 2014.

[54] Fabian Pedregosa, Gaël Varoquaux, Alexandre Gramfort, Vincent Michel, Bertrand Thirion, Olivier Grisel, Mathieu Blondel, Peter Prettenhofer, Ron Weiss, Vincent Dubourg, et al. Scikitlearn: Machine learning in Python. the Journal of machine Learning research, 12:2825–2830, 2011.

[55] Ruopeng Xie, Jiahui Li, Jiawei Wang, Wei Dai, André Leier Tatiana T Marquez-Lago, Tatsuya Akutsu, Trevor Lithgow, Jiangning Song, and Yanju Zhang. DeepVF: a deep learning-based hybrid framework for identifying virulence factors using the stacking strategy. Briefings in bioinformatics, 22(3):bbaa125, 2021.

[56] Jorge R Vergara and Pablo A Estévez. A review of feature selection methods based on mutual information. Neural computing and applications, 24:175–186, 2014.

[57] Chun-Qiu Xia, Xiaoyong Pan, and Hong-Bin Shen. Protein–ligand binding residue prediction enhancement through hybrid deep heterogeneous learning of sequence and structure data. Bioinformatics, 36(10):3018–3027, 2020.

[58] Laurens Van der Maaten and Geoffrey Hinton. Visualizing data using t-SNE. Journal of machine learning research, 9(11), 2008.

[59] Subash C Pakhrin, Suresh Pokharel, Kiyoko F Aoki-Kinoshita, Moriah R Beck, Tarun K Dam, Doina Caragea, and Dukka B Kc. LMNglyPred: prediction of human N-linked glycosylation sites using embeddings from a pre-trained protein language model. Glycobiology, 33(5):411–422, 2023.

[60] Alhasan Alkuhlani, Walaa Gad, Mohamed Roushdy, and Abdel-Badeeh M Salem. Pustackngly: positive-unlabeled and stacking learning for n-linked glycosylation site prediction. IEEE Access, 10:12702–12713, 2022.

[61] Gul Rukh, Shahid Akbar, Gauhar Rehman, Fawaz Khaled Alarfaj, and Quan Zou. StackedEnC-AOP: prediction of antioxidant proteins using transform evolutionary and sequential features based multi-scale vector with stacked ensemble learning. BMC bioinformatics, 25(1):256, 2024.

[62] Islam Uddin, Hamid Hussain Awan, Majdi Khalid, Salman Khan, Shahid Akbar, Mahidur R Sarker, Maher GM Abdolrasol, and Thamer AH Alghamdi. A hybrid residue based sequential encoding mechanism with XGBoost improved ensemble model for identifying 5-hydroxymethylcytosine modifications. Scientific Reports, 14(1):20819, 2024.

[63] Guangchen Liu, Xun Chen, Yihui Luan, and Dawei Li. Viruspredictor: Xgboost-based software to predict virus-related sequences in human data. Bioinformatics, 40(4):btae192, 2024.

[64] Thi-Oanh Tran and Nguyen Quoc Khanh Le. Sa-ttca: an svm-based approach for tumor t-cell antigen classification using features extracted from biological sequencing and natural language processing. Computers in Biology and Medicine, 174:108408, 2024.

[65] Arvind Kumar Yadav, Pradeep Kumar Gupta, and Tiratha Raj Singh. PMTPred: machinelearning-based prediction of protein methyltransferases using the composition of k-spaced amino acid pairs. Molecular Diversity, 28(4):2301–2315, 2024.

[66] Ian Goodfellow. Deep learning, 2016.

[67] John Jumper, Richard Evans, Alexander Pritzel, Tim Green, Michael Figurnov, Olaf Ronneberger, Kathryn Tunyasuvunakool, Russ Bates, Augustin Žídek, Anna Potapenko, et al. Highly accurate protein structure prediction with AlphaFold. nature, 596(7873):583–589, 2021.

